# FoxP3 recognizes microsatellites and bridges DNA through multimerization

**DOI:** 10.1101/2023.07.12.548762

**Authors:** Wenxiang Zhang, Fangwei Leng, Xi Wang, Ricardo N. Ramirez, Jinseok Park, Christophe Benoist, Sun Hur

**Author notes:** These authors contributed equally (Wenxiang Zhang, Fangwei Leng).

## Abstract

FoxP3 is a transcription factor (TF) essential for development of regulatory T cells (Tregs), a branch of T cells that suppress excessive inflammation and autoimmunity^1-5^. Molecular mechanisms of FoxP3, however, remain elusive. We here show that FoxP3 utilizes the Forkhead domain––a DNA binding domain (DBD) that is commonly thought to function as a monomer or dimer––to form a higher-order multimer upon binding to T_n_G repeat microsatellites. A cryo-electron microscopy structure of FoxP3 in complex with T_3_G repeats reveals a ladder-like architecture, where two double-stranded DNA molecules form the two “side rails” bridged by five pairs of FoxP3 molecules, with each pair forming a “rung”. Each FoxP3 subunit occupies TGTTTGT within the repeats in the manner indistinguishable from that of FoxP3 bound to the Forkhead consensus motif (FKHM; TGTTTAC). Mutations in the “intra-rung” interface impair T_n_G repeat recognition, DNA bridging and cellular functions of FoxP3, all without affecting FKHM binding. FoxP3 can tolerate variable “inter-rung” spacings, explaining its broad specificity for T_n_G repeat-like sequences *in vivo* and *in vitro*. Both FoxP3 orthologs and paralogs show similar T_n_G repeat recognition and DNA bridging. These findings thus reveal a new mode of DNA recognition that involves TF homo-multimerization and DNA bridging, and further implicates microsatellites in transcriptional regulation and diseases.

## Introduction

How TFs utilize a limited repertoire of DBDs to orchestrate complex gene regulatory networks is a central and yet unresolved question^6-9^. While certain TFs, such as those with zinc-finger (ZF) DBDs, can expand the complexity of their sequence specificity by forming an array of DBDs, the vast majority of TFs utilize a single DBD with narrow sequence specificity shared with other members of the DBD family^7^. One prominent model to rationalize this apparent paradox is that cooperative actions of multiple distinct TFs with distinct DBDs give rise to combinatorial complexity^10,11^. However, whether a single TF with a single DBD can also recognize distinct sequences on its own and perform divergent transcriptional functions, depending on the conformation or multimerization state, has not been fully addressed.

FoxP3 is an essential TF in Tregs development, of which loss-of-function mutations cause a severe multiorgan autoimmune disease, immune dysregulation, <u>p</u>olyendocrinopathy, <u>e</u>nteropathy, <u>X</u>-linked (IPEX) syndrome^1-5^. Previous studies showed that FoxP3 remodels the global transcriptome and three-dimensional genome organization in the late stage of Treg development^12-15^. However, molecular mechanisms of FoxP3, including its direct target genes and *in vivo* sequence specificity, remain unclear^13-16^.

FoxP3 DNA binding is primarily mediated by a Forkhead domain, which is shared among ∼50 TFs of the Forkhead family^17,18^. Most Forkhead domains form a conserved winged-helix conformation and recognize the consensus FKHM sequence (TGTTTAC)^19^. While isolated Forkhead domain of FoxP3 was originally crystallized as an unusual domain-swap dimer^20,21^, a recent study showed that FoxP3 does not form a domain-swap dimer but instead folds into the canonical winged-helix conformation in the presence of the adjacent Runx1-binding region (RBR)^22^. It was further shown that FoxP3 has a strong preference for inverted repeat FKHM (IR-FKHM) over a single FKHM *in vitro* by forming a head-to-head dimer^22^. However, previous chromatin immunoprecipitation sequencing (ChIP-seq)^14,23,24^ and Cleavage Under Targets & Release Using Nuclease sequencing (CNR-seq)^14^ analyses did not reveal enrichment of IR-FKHM in FoxP3-occupied genomic regions within cells^22^. While individual FKHM is present in ∼10% of the FoxP3 ChIP peaks, they too may not be the FoxP3 binding sites as DNase I protection patterns at these sites were unaffected by FoxP3 deletion^24^. These observations raised the question of what sequences FoxP3 in fact recognizes in cells and whether FoxP3 can employ a previously unknown mode of binding to recognize new sequence motifs distinct from FKHM.

## Results

### FoxP3 binds T_n_G repeat microsatellites

To re-evaluate FoxP3 sequence specificity, we performed an unbiased pull-down of genomic DNA with recombinant FoxP3 protein. Use of genomic DNA, as opposed to synthetic DNA oligos, enables testing sequence specificity in the context of naturally existing repertoire of sequences. It can also allow identification of longer motifs by using genomic DNA fragments longer than ∼20-40 bp––the typical lengths used in previous biochemical studies of FoxP3^22,25,26^. We isolated genomic DNA from mouse EL4 cells, fragmented to ∼100-200 bp, incubated with purified, MBP-tagged mouse FoxP3 and performed MBP pull-down, followed by next-generation sequencing (NGS) of the co-purified DNA (FoxP3 PD-seq, Fig. 1a). We used recombinant FoxP3 protein (FoxP3^ΔN^) harboring zinc finger (ZF), coiled coil (CC), RBR and Forkhead domains but lacking the N-terminal Pro-rich region (Fig. 1a). FoxP3^ΔN^ was previously shown to display the same DNA specificity as full-length FoxP3 among the test set^22^. *De novo* motif analysis showed a strong enrichment of T_n_G repeats (n=2-5) by FoxP3 pull-down, using either MBP alone pull-down or input as a control (Fig. 1b, Supplementary Table 1a). T_3_G repeat sequence was the highest-ranking motif, which accounted for 49.8% of the peaks. No other motifs, including the canonical FKHM or other repeats, were similarly enriched (Supplementary Table 1a). FoxP3 pull-down using nucleosomal DNA from EL4 cells revealed similar enrichment of T_n_G repeat-like sequences (Supplementary Table 1a).

**Figure 1.**
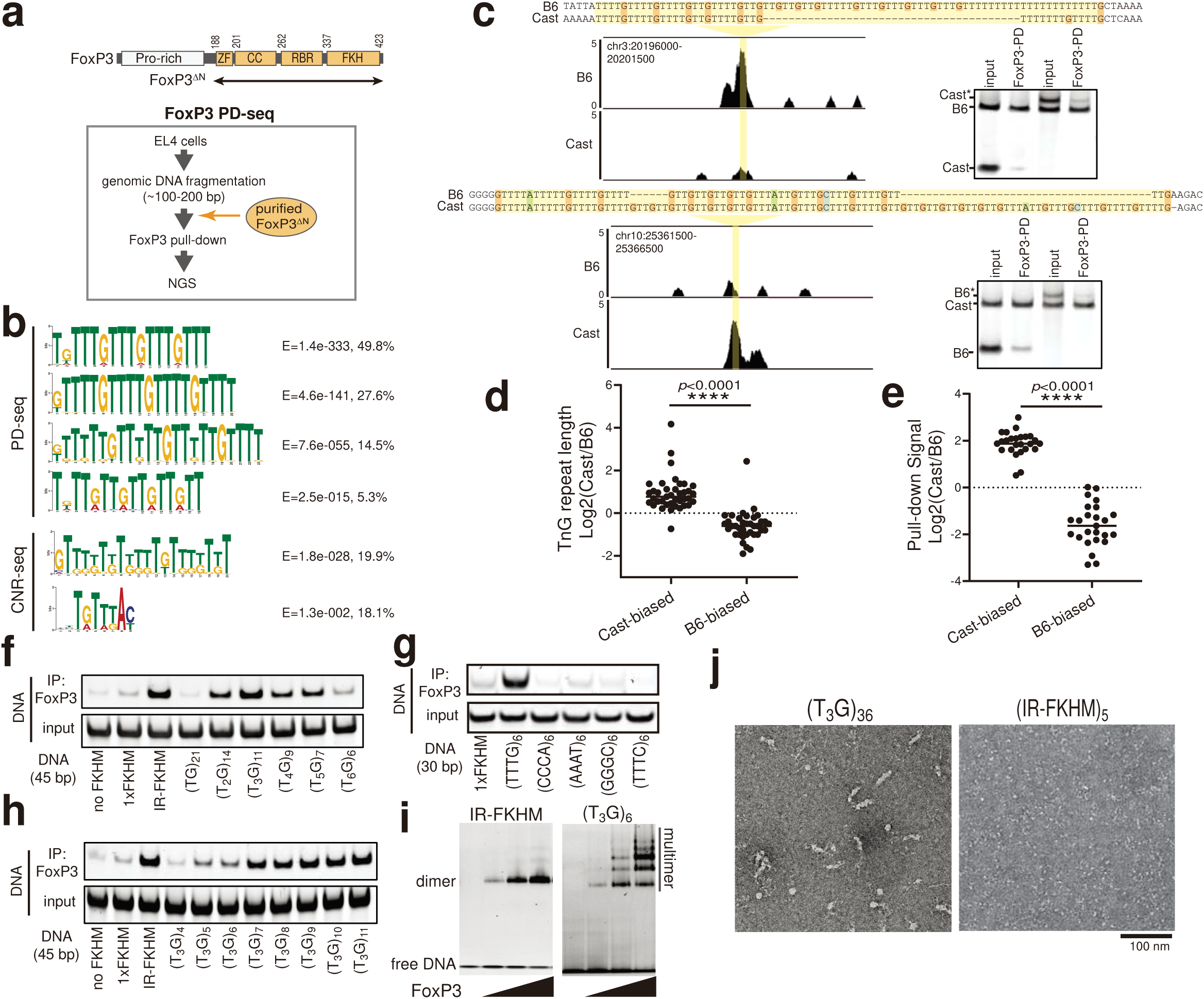
FoxP3 recognizes T_n_G repeat microsatellites. a. FoxP3 domain architecture and schematic of FoxP3 pull-down (PD)-seq. ZF: zinc finger; CC: coiled-coil; RBR: Runx1-binding region. b. *De novo* motif analysis of FoxP3 PD-seq peaks (n=21,605) and CNR-seq peaks^14^ (n=6,655) by MEME-ChIP and STREME. E-score and percentage of peaks containing the given motif are shown on the right. See also Supplementary Table 1a and 1b for the comprehensive list of motifs for PD-seq, CNR-seq^12,14^ and ChIP-seq data^14,23^. c. Allelic imbalance in FoxP3 binding *in vivo*. Left: genome browser view of CNR-seq^14^ showing B6-biased (top) and Cast-biased (bottom) peaks. B6 genomic coordinates are shown on the upper left corner. Right: B6 and Cast DNA oligos were mixed 1:1 and subjected to FoxP3^ΔN^ pull-down and gel analysis. Cast* and B6* represent oligos extended with the random sequence (Supplementary Table 2b) to reverse their length bias. d. T_n_G repeat-length comparison between Cast and B6 at 78 loci showing allelic bias in (c). Repeat lengths were measured in nucleotide. n = 25 for Cast-biased loci and n = 25 for B6-biased loci were used for this comparison. Two-tailed unpaired t-tests. ****, *p* < 0.0001. e. Allelic imbalance in FoxP3 binding *in vitro*. 50 pairs of Cast and B6 sequences (Supplementary Table 2a) were chosen from the 78 pairs in (d) and subjected to FoxP3^ΔN^ pull-down. For each pair, the recovery rate of the Cast and B6 DNA were measured and their ratios were plotted. Each data point represents an average of the two pull-downs. Two-tailed unpaired t-tests. ****, *p* < 0.0001. f-h. FoxP3–DNA interaction as measured by FoxP3^ΔN^ pull-down. DNA harboring a random sequence (no FKHM), a single FKHM (1XFKHM), IR-FKHM or tandem repeats of T_n_G (n=1-6) were used. All DNAs were 45 bp-long in (f) and (h), and 30 bp-long in (g). i. Native gel shift assay of MBP-tagged FoxP3^ΔN^ (0-0.4 μM) with DNA (30 bp, 0.05 μM) containing IR-FKHM or (T_3_G)_6_. j. Representative negative-stain EM images of FoxP3^ΔN^ in complex with (T_3_G)_36_ and (IR-FKHM)_5_. Both DNAs were 144 bp-long.

*De novo* motif analysis of previously published FoxP3 CNR-seq^12,14^ and ChIP-seq data^14,23^ also identified T_n_G repeat-like motifs as one of the most significant motifs in all four datasets (Fig. 1b, Supplementary Table 1b). The enrichment score for T_n_G repeat-like motifs (E-value) was more significant than that of FKHM in all cases (Fig. 1b, Supplementary Table 1b). Note that T_n_G repeat-like motifs have not been reported from these original studies, likely reflecting the common practice of discarding simple repeats in motif analysis. T_n_G repeat-like motifs were not identified from open chromatin regions (as measured by ATAC-seq^27^) in Treg cells that were not occupied by FoxP3 (Supplementary Table 1b).

To examine whether T_n_G repeat-like sequences indeed contribute to FoxP3–DNA interaction in Treg cells, we analyzed published FoxP3 CNR-seq data generated using F1 hybrids of B6 and Cast mice strains^14^. Because of the wide divergence between B6 and Cast genomes, such data allow evaluation of the impact of sequence variations on TF binding. Out of 196 sites showing allelic imbalance (fold change >=4) in FoxP3 CNR-seq, 78 sites harbored T_n_G repeat-like elements in at least one allele, the frequency (39.3%) significantly higher than that in the mouse genome (∼0.06%, *p*<1e-8, Extended Data Fig. 1a). Furthermore, all but 4 sites showed T_n_G repeat length mirroring the allelic bias in FoxP3 occupancy (Fig. 1c, 1d). Of the 78 sites, we randomly chose 50 sites, 25 each from B6- and Cast-biased peaks, and tested FoxP3 binding efficiency by FoxP3^ΔN^ pull-down. Out of the 50 pairs of sequences tested, pull-down efficiency of 47 pairs recapitulated differential binding in CNR-seq (Figs. 1c, 1e). All 47 sites showed significant truncations in the T_n_G repeats in the less-preferred allele (see Supplementary Table 2a for the full list of sequences). Note that the pull-down preference for longer T_n_G repeats was not due to the different DNA lengths used––an extension of the less-preferred allele sequences with a random sequence at a DNA end (B6* and Cast* in Fig. 1c, see Supplementary Table 2b for sequence) did not alter the allele bias. Altogether, these results suggest that T_n_G repeat-like elements play an important role in FoxP3–DNA interaction *in vitro* and *in vivo*.

Genome-wide analysis showed that there are 46,574 loci in the *M. musculus* genome with T_n_G repeat-like sequences and that they are predominantly located distal to annotated transcription start sites (TSSs) with 9.5% residing within 3 kb of the annotated TSSs (Extended Data Figs. 1a,1b). In comparison, among the T_n_G repeat-containing FoxP3 CNR peaks^12,14^ (n=3,301 out of the 9,062 CNR peaks), 38.4% were found within 3 kb of TSSs (Extended Data Fig. 1c). T_n_G repeat-containing FoxP3 CNR peaks also displayed higher levels of trimethylated H3K4 (H3K4me3), acetylated H3K27 (H3K27ac) and chromatin accessibility than the genome-wide T_n_G repeats (Extended Data Figs. 1d, 1e, 1f). These results suggest that while T_n_G repeat-like sequences are common in the *M. musculus* genome, FoxP3 utilizes a small fraction of T_n_G repeat-like sequences in accessible, functional sites for transcriptional regulation.

### FoxP3 multimerizes on T_n_G repeats

To examine T_n_G repeat enrichment in PD-seq and CNR/ChIP-seq indeed represent previously unrecognized sequence specificity of FoxP3, we compared FoxP3 binding to DNA with T_n_G repeats (n=1 to 6) vs. those containing IR-FKHM, the highest affinity sequence reported for FoxP3 to date^22^. All DNAs were 45 bp-long (see Supplementary Table 2b for sequences). We found that T_3_G repeat was comparable to IR-FKHM in FoxP3 binding and was the tightest binder among the T_n_G repeats (Fig. 1f), consistent with it being the most significant motif in PD-seq (Fig. 1b). T_2_G, T_4_G and T_5_G repeats also showed more efficient binding than a single FKHM (1XFKHM) or random sequence (no FKHM). No other simple repeats showed FoxP3 binding comparable to T_3_G repeats (Fig. 1g). FoxP3 affinity increased with the copy number of T_3_G, when compared among DNAs of the same length (Fig. 1h). The preference for T_3_G repeats was also seen using full-length FoxP3 expressed in HEK293T cells (Extended Data Fig. 1g) or when pull-down bait was switched from FoxP3 to DNA (Extended Data Fig. 1h). Finally, FoxP3 can bind T_3_G repeats even in the presence of nucleosomes (Extended Data Fig. 1i), suggesting that similar interaction can occur in the context of chromatinized DNA.

We next investigated how FoxP3 recognizes T_3_G repeats. Unlike IR-FKHM, T_3_G-repeat DNA induced FoxP3 multimerization as indicated by slowly migrating species in native gel-shift assay (Fig. 1i). Protein-protein crosslinking also suggested higher-order multimerization in the presence of T_3_G repeats, but not with IR-FKHM or 1XFKHM (Extended Data Fig. 1j). In support of T_3_G repeat-induced multimerization, MBP-tagged FoxP3 was co-purified with GST-tagged FoxP3 only in the presence of T_3_G repeat, but not with IR-FKHM (Extended Data Fig. 1k). Finally, negative electron microscopy revealed a filamentous multimeric architecture of FoxP3 on 36 tandem repeats of T_3_G (Fig. 1j), the copy number chosen to aid clear visualization. Other DNAs of the same length, such as (A_3_G)_36_, (TGTG)_36_ or (IR-FKHM)_5_, did not show similar multimeric architectures under equivalent conditions (Fig. 1j, Extended Data Fig. 1l). These results suggest that FoxP3 forms distinct multimers on T_3_G repeats.

### Structure of FoxP3 bound to T_3_G repeats

To understand how FoxP3 forms multimers on T_3_G repeats, we determined the cryo-electron microscopy (cryo-EM) structure of FoxP3^ΔN^ in complex with (T_3_G)_18_. Single particle reconstruction led to a 3.6 Å resolution map after global refinement and a 3.3 Å resolution map after focused refinement of the central region (Extended Data Figs. 2a-2f, Extended Data Table 1). The density map revealed two continuous double-stranded DNA molecules spanning ∼50 bp (Fig. 2a). Both DNA molecules adopted the classic B-form DNA with the average twist angle of 33.5°/bp and the average rise of 3.19 Å/bp. The density map could also be fitted with the crystal structure of DNA-bound FoxP3 monomer containing part of RBR and Forkhead (residue 326-412), allowing placement of ten FoxP3 subunits without ZF, CC and RBR residues 188-325. Only the non-swap, winged-helix conformation was compatible with the density map (Extended Data Fig. 2g). Consistent with this, R337Q, a loss-of-function IPEX mutation that induces domain-swap dimerization^22^, showed significantly reduced affinity for T_3_G repeats (Extended Data Fig. 2h).

**Figure 2.**
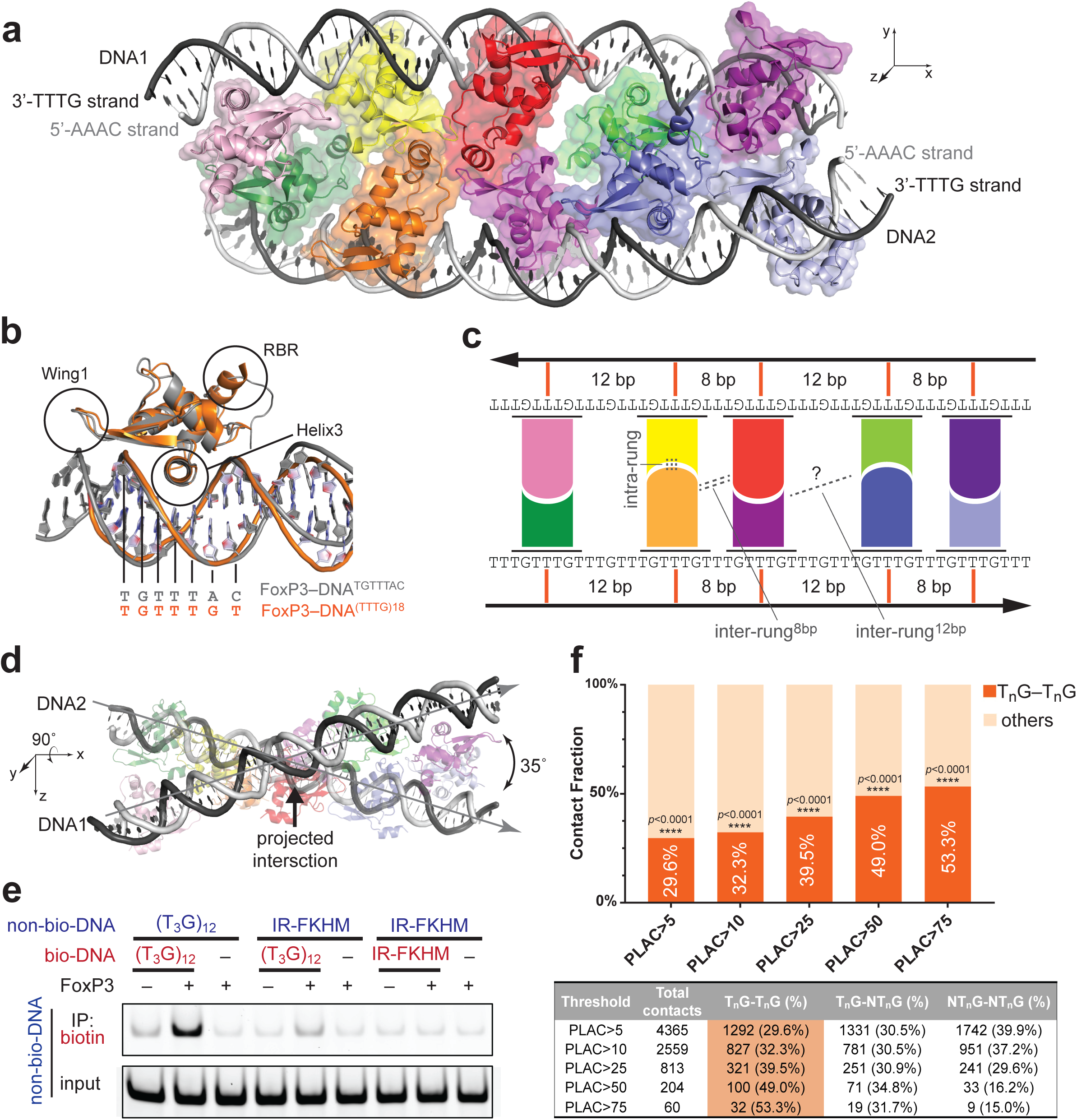
FoxP3 forms a ladder-like multimer upon binding to T_3_G repeat DNA. a. Cryo-EM structure of FoxP3^ΔN^ decamer in complex with two DNA molecules (grey) harboring (T_3_G)_18_. Each of the ten FoxP3 subunits are colored differently. b. Comparison of a representative FoxP3^ΔN^ subunit from (a) (orange) with a FoxP3^ΔN^ subunit from the head-to-head dimeric structure (grey, PDB: 7TDX). Helix3 recognizes DNA sequence (TGTTTAC in the head-to-head dimer, TGTTTGT in the ladder-like multimer) by inserting it into the major groove. c. Schematic of the ladder-like architecture of FoxP3 on T_3_G repeat DNA. d. Skew relationship between the two DNA molecules, which is evident when looking down the y-axis of (a). e. DNA bridging assay. Biotinylated DNA (bio-DNA, 82 bp) and non-biotinylated DNA (non-bio-DNA, 60 bp) were mixed at 1:1 ratio (0.1 μM each), incubated with FoxP3^ΔN^ (0.4 μM), and subjected to streptavidin pull-down prior to gel analysis. Non-biotinylated DNA in the eluate was visualized by SybrGold staining. f. Chromatin contacts at FoxP3-bound anchors identified from HiC- and PLAC-seq^12^. Contacts with frequency>5 in WT Treg HiC and connected by two FoxP3-bound anchors were analyzed with an increasing FoxP3 PLAC-seq count threshold. Percentage of the unique contacts mediated by two T_n_G anchors (out of all unique contacts between two FoxP3-bound anchors) was indicated. All T_n_G-T_n_G contacts were between two distinct 10 kb anchor bins. **** indicates *p*<0.0001 using the null hypothesis that the T_n_G-T_n_G contacts occur by random chance.

DNA sequence assignment (see Extended Data Fig. 3a) revealed that all ten FoxP3 subunits interacted with T_3_G repeat DNA in the manner indistinguishable from that of FoxP3 bound to the canonical FKHM, recognizing TGTTTGT in place of TGTTTAC (Fig. 2b). This FoxP3-DNA register was further confirmed by FoxP3 footprint analysis using DNA mutagenesis and NFAT– FoxP3 cooperativity (Extended Data Figs. 3b, 3c). Note that NFAT is a known interaction partner of FoxP3 and assists FoxP3 binding to DNA only when their binding sites are 3 bp apart, the property used for inferring FoxP3–DNA registers (see Extended Data Fig. 3b for details).

The overall architecture resembled a ladder where the two double-stranded DNA molecules formed side rails bridged by five “rungs”, each of which consisted of two FoxP3 subunits bound to different DNA and joined by direct protein-protein interactions (“intra-rung” interactions) (Fig. 2a, 2c). These rungs were separated by 8 bp or 12 bp in an alternating fashion, forming two different types of “inter-rung” interactions (inter-rung^8bp^ and inter-rung^12bp^) with divergent significance (to be discussed later). Given that both DNA molecules had the helical periodicity of 10.7 bp per turn, this alternating spacing pattern allowed FoxP3 molecules to occupy consecutive major grooves on one side of each DNA. This geometry, in turn, enabled the FoxP3 molecules on opposing DNA to face each other and form the “rungs” of the ladder. None of the intra- and inter-rung interactions resembled the previously reported head-to-head dimerization interaction (Extended Data Fig. 3d)^22^, revealing a new mode of molecular assembly for FoxP3.

The two DNA molecules are skew to each other (i.e. non-parallel, non-intersecting). When projected onto the x-y plane as in Fig. 2a, they appeared parallel, but projection onto the x-z plane as in Fig. 2d suggested that they approached each other at an angle of 35°. The divergence of the two DNA molecules can explain why the multimeric assembly was limited to the decamer spanning ∼50 bp near the projected intersection point (Fig. 2d), even though the DNA sample in cryo-EM was 72 bp-long and had many more T_3_G repeats to accommodate additional FoxP3 molecules. The lack of cryo-EM density for FoxP3 molecules bound to DNA without forming the “rung” suggests that the intra-rung interaction is critical for stable protein-DNA interaction. In other words, DNA bridging may be an integral part of the assembly.

To test whether DNA bridging indeed occurs in solution, we examined co-purification of non-biotinylated DNA (prey) with biotinylated DNA (bait) in the presence and absence of FoxP3. We observed DNA bridging between biotinylated and non-biotinylated T_3_G repeats only in the presence of FoxP3^ΔN^ (Fig. 2e). DNA bridging was not observed between IR-FKHM and IR-FKHM DNAs or between (T_3_G)_12_ and IR-FKHM DNAs. Similar T_n_G repeat-dependent bridging was observed with full-length FoxP3 expressed in 293T cells (Extended Data Fig. 3e). Additionally, T_3_G repeat DNA bridging occurred more efficiently with an increasing concentration of FoxP3 (Extended Data Fig. 3f), suggesting that DNA bridging is not an artificial consequence of saturating multimeric FoxP3 with DNA.

To further examine whether FoxP3 binding to T_n_G repeats mediates long-distance chromatin contacts in Tregs, we analyzed the available HiC-seq, PLAC-seq and HiChIP-seq data^12,13^. The limited resolution of these data (5-10kb) precluded direct motif analysis of the chromatin contact anchors. Instead, we asked how frequently contacts are made between anchors containing FoxP3 CNR peaks with T_n_G repeats (T_n_G anchors) vs. those lacking T_n_G repeats (non-T_n_G anchors). Among the high-frequency contacts (HiC frequency>5, PLAC frequency>5-75) between FoxP3-bound anchors, we found that those between two T_n_G anchors (30-53%) were more prevalent than expected by chance (13.7%) and that such T_n_G–T_n_G contacts were more enriched among the stronger contacts (Fig. 2f, Supplementary Table 3, tabs 1-6). In contrast, non-T_n_G–non-T_n_G contacts were more depleted among the stronger contacts. This is despite the fact that non-T_n_G CNR peaks have higher levels of chromatin accessibility and H3K4me3 than T_n_G CNR peaks, while displaying similar H3K27ac levels (Extended Data Figs. 4a-4c). Most of the T_n_G–T_n_G contacts showed increased frequency in WT Tregs relative to in FoxP3 knock-out Treg-like cells (Extended Data Fig. 4d). Furthermore, many of the anchors for the T_n_G–T_n_G contacts were nearby Treg signature genes (e.g. *Il2ra, Cd28, Tnfaip3, Ets1* in Supplementary Table 3, tab 7), and overlapped with previously characterized enhancer-promoter loop anchors in Tregs (Extended Data Fig. 4e), implicating their transcriptional functions. These results together support that FoxP3 multimerization on T_n_G repeats contributes to long-distance chromatin contacts in Tregs.

### Intra-rung interaction is essential

Examination of the intra-rung interaction showed that multiple distinct parts of the protein are involved; wing1 (W1), a loop between helix 2 and 4 (H2/H4 loop) and Helix6 (H6) of one subunit interacted with RBR and H2/H4 of the other subunit within the rung (Fig. 3a). While the resolution at the interface was insufficient to assign precise side chain conformations, the structure identified R356 in the H2/H4 loop, V396 and V398 in W1, D409/E410/F411 in H6 as residues at the interface (Fig. 3a). We also chose V408 in H6, which was adjacent to the interface residues and is mutated to Met in a subset of IPEX patients^15,28,29^. Mutations of these interface residues, including V408M, disrupted T_3_G repeat binding (Fig. 3b, right) and DNA bridging (Fig. 3c). The same mutations had a minimal impact on IR-FKHM binding (Fig. 3b, left), which requires head-to-head dimerization of FoxP3^22^. This is consistent with the previous structure showing that these residues are far from either the DNA binding or the head-to-head dimerization interface^22^. The negative effect of the intra-rung mutations on T_3_G repeat binding as well as DNA bridging further supports that DNA bridging is required for FoxP3 multimerization on T_3_G repeats, rather than a simple consequence of FoxP3 multimerization.

**Figure 3.**
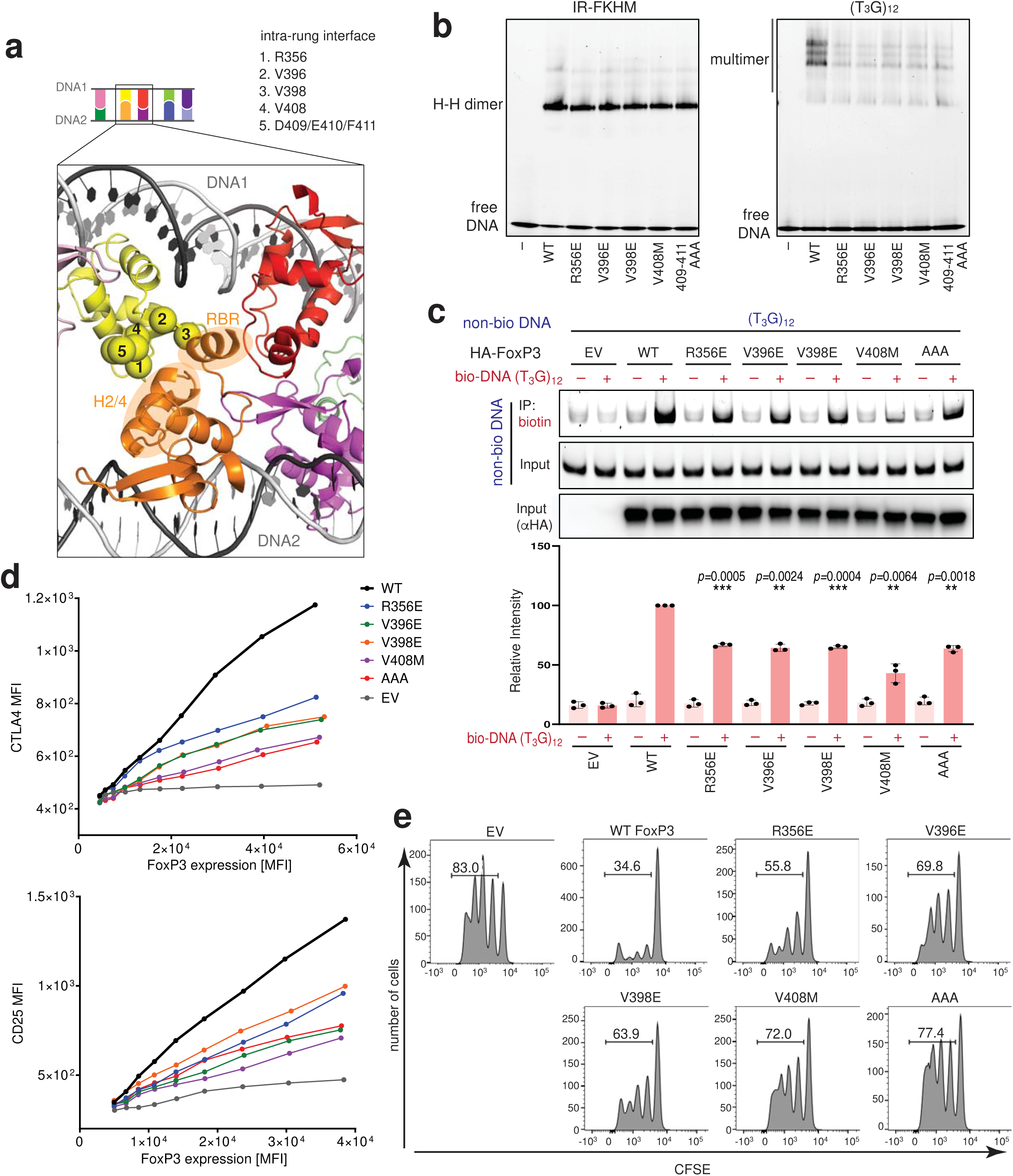
Intra-rung interaction is essential for T_n_G repeat recognition, DNA bridging, and cellular functions of FoxP3. a. Intra-rung interface. Ca of R356, V396, V398, V408 and D409/E410/F411 are shown in spheres. These residues on yellow subunit interact with RBR and helix2-helix4 (H2/4) of the orange subunit. Subunit colors are as in Fig. 2a. b. Effect of intra-rung interface mutations on DNA binding. MBP-tagged FoxP3^ΔN^ (0.4 μM) was incubated with IR-FKHM or (T_3_G)_12_ (60 bp for both) and analyzed by native gel shift assay. c. Effect of intra-rung interface mutations on DNA bridging. FoxP3 (or empty vector, EV) was expressed in 293T cells and lysate was incubated with a mixture of biotinylated and non-biotinylated DNA (1:1 ratio), followed by streptavidin pull-down and gel analysis. Relative level of non-biotinylated DNA co-purified with biotinylated DNA was quantitated from three independent pull-downs. The difference was compared with WT in the presence of bio-DNA. Two-tailed paired t-tests. *p*<0.001 for *** and *p*<0.005 for **. d. Transcriptional activity of FoxP3. CD4+ T cells were retrovirally transduced to express FoxP3, and its transcriptional activity was analyzed by measuring the level of the known target genes CTLA4 and CD25 using FACS. FoxP3 level was measured using Thy 1.1, which is under the control of IRES from the bicistronic mRNA for FoxP3. MFI indicates mean fluorescence intensity. e. T cell suppression assay of intra-rung interface mutations. FoxP3-transduced T cells (suppressors) were mixed with naïve T cells (responders) at a 1:2 ratio and the effect of the suppressor cells on proliferation of the responder cells were measured by the CFSE dilution profile of the responder T cells.

These intra-rung mutations disrupted cellular functions of FoxP3, as measured by FoxP3-induced gene expression (*e.g. CTLA4* and *CD25* protein levels in Fig 3d and RNA-seq in Extended Data Fig. 5a), target loci binding (as measured by FoxP3 ChIP-seq in Extended Data Fig. 5b) and T cell-suppressive functions (Fig. 3e). None of these mutations affected nuclear localization, the level of FoxP3 (Extended Data Figs. 5c, 5d) or FoxP3’s interaction with NFAT (Extended Data Fig. 5e), although slight reduction in NFAT binding was seen for V398E. These results suggest that the ladder-like assembly is important for the transcriptional functions of FoxP3.

### Relaxed sequence specificity of multimer

We next examined the potential role of the inter-rung interactions. The inter-rung^8bp^ interaction was mediated by RBR–RBR contacts, which displayed continuous EM density indicative of a strong or ordered interaction (Extended Data Fig. 2f, Fig. 4a). Unlike the intra-rung interface mutations, mutations in RBR, for example F331D, disrupted FoxP3 binding to both T_3_G repeats and IR-FKHM^22^ (Fig. 4b, Extended Data Fig. 6a), suggesting that RBR plays an important role in both ladder-like multimerization and head-to-head dimerization^22^. Consistent with the importance of the inter-rung^8bp^ interaction, changes in the inter-rung^8bp^ spacing from 8 bp (1 bp gap) to 9 bp (2 bp gap) or 7 bp (no gap) resulted in a significant impairment in FoxP3 binding to T_3_G repeats (Fig. 4c).

**Figure 4.**
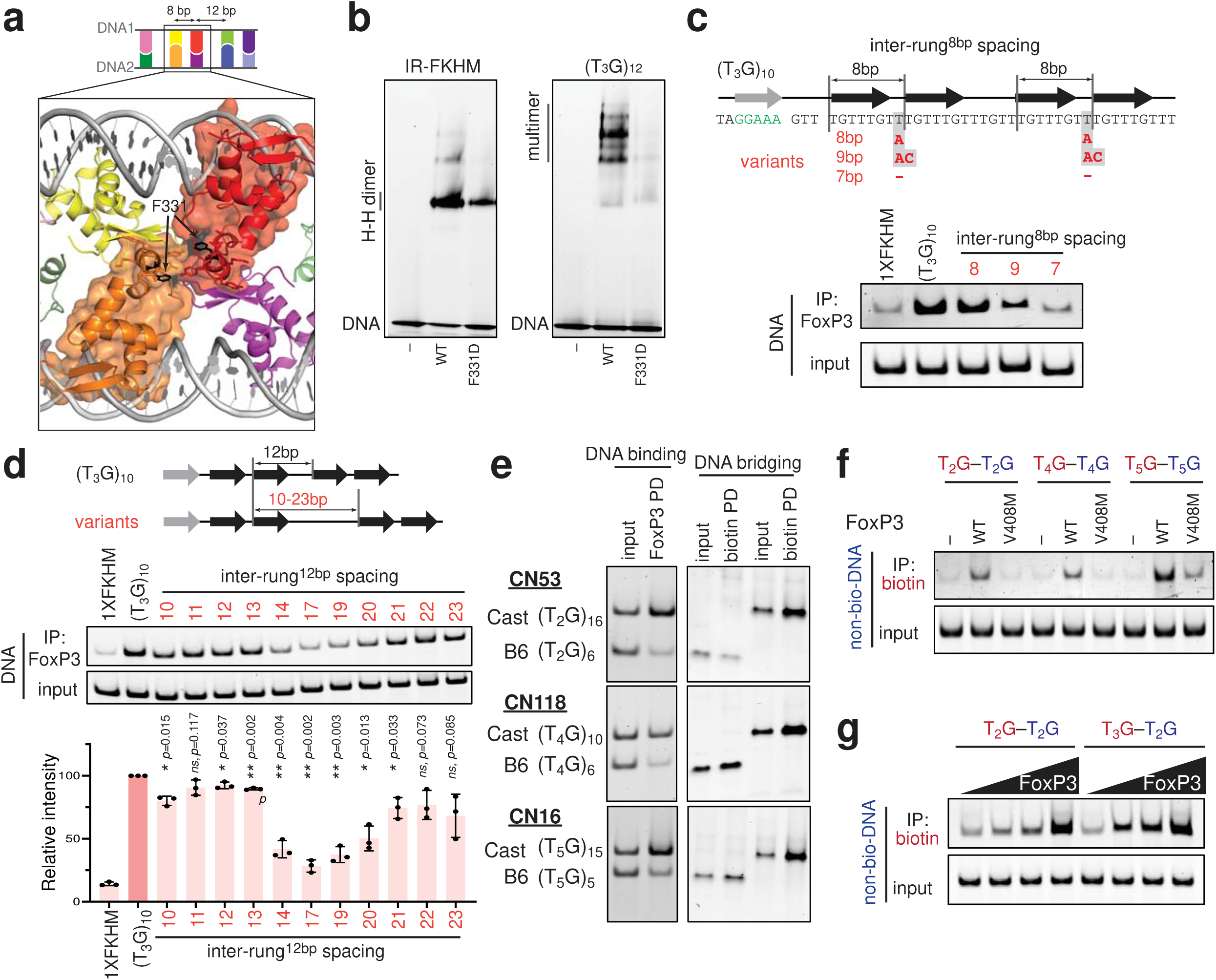
Architectural flexibility of the ladder-like assembly broadens sequence specificity of FoxP3. a. Structure highlighting the inter-rung^8bp^ interaction between orange and red subunits (in surface representation). The interaction is primarily between RBRs, where F331 resides. b. Effect of inter-rung^8bp^ mutation (F331D) on FoxP3–DNA interaction by FoxP3^ΔN^ pull-down. c. Effect of inter-rung^8bp^ spacings on FoxP3 binding by FoxP3^ΔN^ pull-down. Both inter-rung^8bp^ gap nucleotides were changed from T in (T_3_G)_10_ to A (8-bp spacing), to AC (9-bp spacing) or to no nucleotide (7-bp spacing). Black arrows indicate FoxP3 footprints. Gray arrow indicates the NFAT binding site. NFAT interacts with FoxP3 and helps fixing the FoxP3-DNA register, which was necessary to examine the effect of DNA sequence variations at or between the FoxP3 footprints. d. Effect of inter-rung^12bp^ spacings on FoxP3 binding by FoxP3^ΔN^ pull-down. The inter-rung^12bp^ spacing was changed from 12 bp in (T_3_G)_10_ to 10-23 bp (see Supplementary Table 2b for sequences). Averaged recovery rate of DNA from three independent pull-downs were plotted. Two-tailed paired t-tests in comparison to (T_3_G)_10_. *p*<0.005 for **, *p*<0.05 for * and *p*>0.05 for ns. e. Comparison of Cast and B6 sequences in FoxP3 binding (left) and DNA bridging (right). Three pairs of sequences at loci CN53, CN118 and CN16 with Cast-bias in CNR-seq were compared (Supplementary Table 2a) by FoxP3^ΔN^ pull-down. f. DNA bridging between (T_2_G)_14_ and (T_2_G)_14_, between (T_4_G)_9_ and (T_4_G)_9_, and between (T_5_G)_7_ and (T_5_G)_7_ in the presence of WT FoxP3 or the IPEX mutant V408M. Biotinylated and non-biotinylated DNA were colored red and blue, respectively. g. DNA bridging between (T_2_G)_14_ and (T_2_G)_14_, and between (T_2_G)_14_ and (T_3_G)_11_ by FoxP3 (0-0.4 μM).

In contrast to the inter-rung^8bp^ interaction, cryo-EM density for the inter-rung^12bp^ interaction was difficult to interpret, which could reflect a weak or less-ordered interaction. In keeping with this, FoxP3 binding tolerated a wide range of inter-rung^12bp^ spacings, with equivalent affinity observed for spacings of 11-13 bp (Fig. 4d). Interestingly, while 14-19 bp spacings were not tolerated, DNA with 21-22 bp spacings showed moderate binding. Given that 11-13 bp, 14-19 bp and 21-22 bp spacings would place FoxP3 one, one and a half and two helical turns away from the upstream FoxP3 molecule, respectively, this cyclical pattern suggests that the precise positions of FoxP3 are not essential for multimeric assembly, so far as the DNA sequence allows FoxP3 molecules to line up on one side of DNA and form the “rungs”. Consistent with the idea, DNA bridging activity showed a similar cyclical pattern (Extended Data Fig. 6b).

This architectural flexibility may explain our observations in Fig. 1, which showed that FoxP3 could bind to a broad range of T_n_G repeat-like sequences besides perfect T_3_G repeats. These include tandem repeats of T_2_G, T_4_G, T_5_G and their various mixtures found in the CNR-seq peaks with allelic imbalance (Supplementary Table 2a). To examine whether a similar ladder-like architecture forms with T_n_G repeat-like sequences that are not perfect T_3_G repeats, we used DNA-bridging activity as a measure of the ladder-like assembly. All 47 pairs of the DNA sequences showing allelic bias in FoxP3 binding *in vivo* and *in vitro* displayed the same allelic bias in DNA bridging (Fig. 4e, Supplementary Table 2a). The multimerization-specific IPEX mutation V408 abrogated bridging of T_2_G, T_4_G, T_5_G repeat DNAs (Fig. 4f), suggesting a similar multimeric architecture for FoxP3 regardless of the exact T_n_G repeat sequences. Interestingly, suboptimal T_n_G repeats (n=2, 4, 5) were bridged with T_3_G repeats more efficiently than with themselves (Fig. 4g, Extended Data Fig. 6c), suggesting that having a strong DNA as a bridging partner helps FoxP3 binding to suboptimal sequences. These results reveal yet another layer of complexity that can broaden the sequence specificity of FoxP3.

### T_n_G repeat binding is conserved in FoxPs

Studies above were done with FoxP3 and T_n_G repeat-like elements from *M. musculus*. We next asked whether T_n_G repeat recognition by FoxP3 is conserved in other species besides *M. musculus*. Inspection of T_n_G repeat-like elements in the *H. sapiens* and *D. rerio* genomes revealed 18,164 and 5,517 distinct sites harboring T_n_G repeats (>29 nt), respectively, in comparison to the 46,574 sites in the *M. musculus* genome (Extended Data Fig. 1a). While T_n_G-like repeats are more frequently located distal to TSSs in all three genomes of *H. sapiens*, *M. musculus* and *D. rerio*, greater fractions are located within ∼3 kb of TSS in higher eukaryotes (12.66%, 9.50% and 5.72% for *H. sapiens*, *M. musculus* and *D. rerio*, respectively) (Extended Data Fig. 1b), even though all three species have similar genes-to-genome size ratios (see Table in Extended Data Fig. 1a). This observation suggests that T_n_G repeats may have been coopted for transcriptional functions in higher eukaryotes.

We examined FoxP3 from *H. sapiens*, *O. anatinus* and *D. rerio*. All three FoxP3 orthologs showed preferential binding to T_3_G repeats and IR-FKHM in comparison to a single FKHM or no FKHM (Extended Data Fig. 6d). They also bridged T_3_G repeats (Extended Data Fig. 6e), suggesting a ladder-like assembly similar to that of *M. musculus* FoxP3. This is in keeping with the fact that the key residues for multimerization were broadly conserved or interchanged with similar amino acids in FoxP3 orthologs (Extended Data Fig. 6f). Given that *D. rerio* FoxP3 represents one of the most distant orthologs from mammalian FoxP3, these results suggest that T_n_G repeat recognition and ladder-like assembly may be ancient properties of FoxP3.

Inspection of sequence alignment of Forkhead TFs revealed that the key residues for multimerization are also well-conserved within the FoxP family, but not outside (Fig. 5a). Biochemical analysis of *M. musculus* FoxP1, FoxP2 and FoxP4 in the FoxP family showed that they preferentially bound T_3_G repeats and bridged T_3_G repeat DNA as with FoxP3 (Figs. 5b, 5c and Extended Data Fig. 6g). *De novo* motif analysis of previously published ChIP-seq data showed that T_n_G repeat-like motifs were indeed enriched in FoxP1- and FoxP4-occupied sites (Fig. 5d, the full list and references included in Supplementary Table 1c). This feature was particularly strong for FoxP1 in lymphoma cell lines (SU-DHL-6 and U-2932) and mouse neural stem cells––T_n_G repeat-like motif was the most significant motif, while FKHM ranked far lower (Fig. 5d). However, in VCap and K-562 cell lines, FoxP1 ChIP-seq peaks did not show T_n_G-like elements, although FKHM was identified as one of the most significant motifs in these cells (Supplementary Table 1c). Similar context-dependent enrichment of T_n_G repeat-like elements was seen with FoxP4, although the motif enrichment was not as strong as with FoxP1 or FoxP3 (Fig. 5d, Supplementary Table 1c). In contrast, long (>10 nt) T_n_G repeat-like elements were not identified from any of the 48 distinct sets of ChIP-seq data for FoxA1, FoxM1, FoxJ2, FoxJ3, FoxQ1 and FoxS1, while FKHM ranked as one of the strongest motifs in many (Supplementary Table 1c). These results suggest that preference for T_n_G repeat-like sequence and ladder-like assembly are conserved properties of FoxP3 paralogs and orthologs, but may not be shared among all Forkhead TFs.

**Figure 5.**
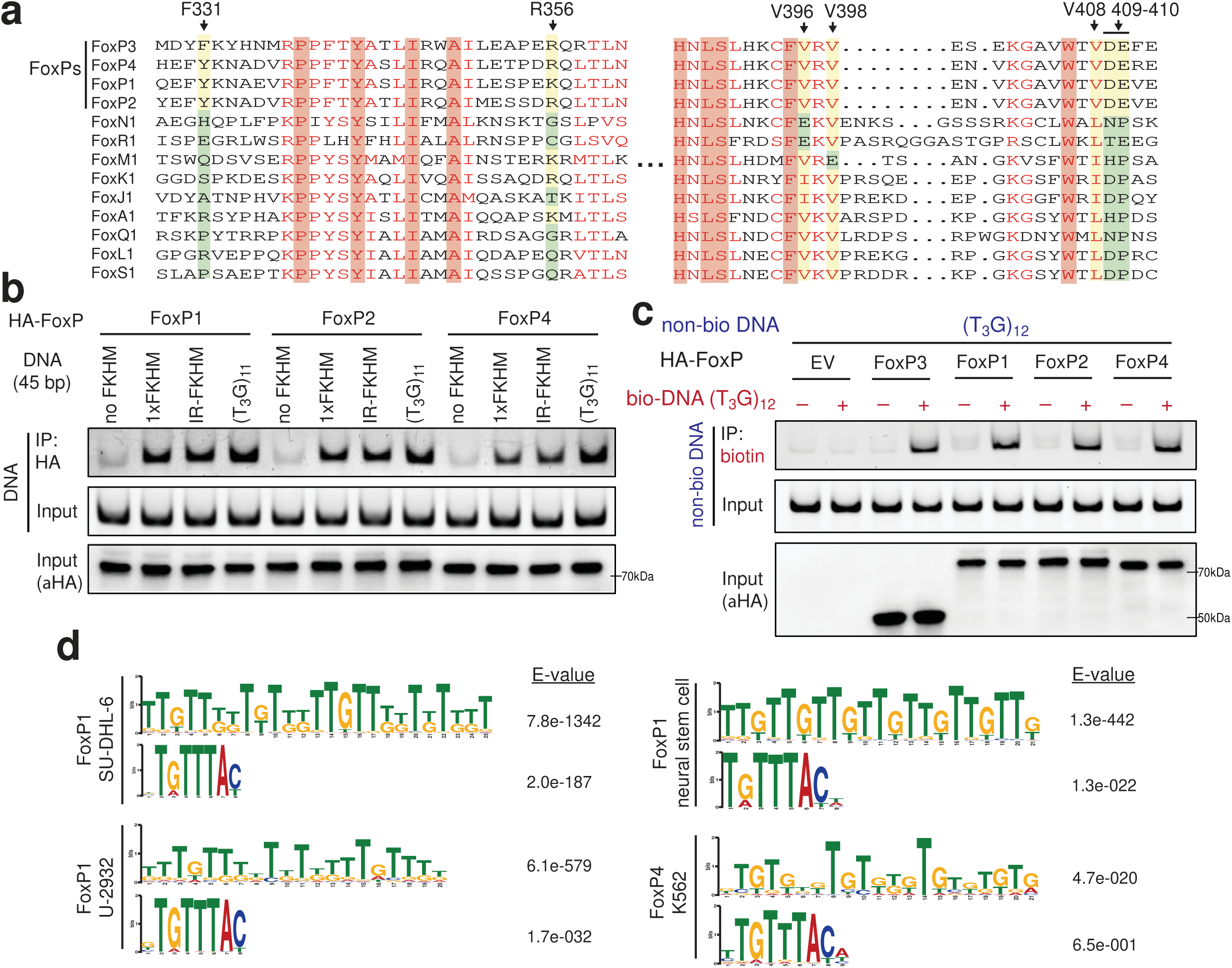
T_n_G microsatellite recognition is conserved among FoxP3 orthologs and paralogs. a. Sequence alignment of forkhead TFs. Residues equivalent to the key interface residues in mouse FoxP3 (arrow on top) were highlighted yellow (when similar to the mouse residues) or green (when dissimilar). b. DNA-binding activity of FoxP1, FoxP2 and FoxP4. HA-tagged FoxP1, FoxP2 and FoxP4 were transiently expressed in HEK293T cells and purified by anti-HA IP. Equivalent amounts of indicated DNAs (all 45 bp) were added to FoxP1/2/4-bound beads and further purified prior to gel analysis (SybrGold). c. DNA-bridging activity of FoxP1, FoxP2 and FoxP4. Experiments were performed as in Fig. 3c using 293T lysate expressing HA-tagged FoxP TFs. d. *De novo* motif analysis of FoxP1 and FoxP4 ChIP-seq peaks from published database ^38,39^. See Supplementary Table 1c for the comprehensive list and their references.

## Discussion

In summary, our findings showed a new mode of TF–DNA interaction that involves TF homo-multimerization and DNA bridging. Upon binding to T_n_G repeats, FoxP3 forms a ladder-like multimer, in which FoxP3 utilizes two DNA molecules as scaffolds to facilitate cooperative multimeric assembly. That is, the first set of FoxP3 molecules (possibly a dimer or two dimers with an 8-bp spacing) that bridge DNA would help recruit additional FoxP3 “rungs”, which would in turn stabilize the bridged DNA architecture and subsequent rounds of FoxP3 recruitment. Such cooperative assembly enables FoxP3 to preferentially target long repeats of T_n_G rather than spurious sequences containing a few copies of T_n_G. The DNA bridging activity also implicates FoxP3 as a previously unrecognized class of TF that can directly mediate architectural functions, which may explain the recently observed role of FoxP3 in chromatin loop formation^12,13^.

How do we reconcile the ladder-like assembly of FoxP3 on T_n_G repeats and the previously reported head-to-head dimeric structure on IR-FKHM or related sequences^22^? Unlike the ladder-like multimerization, cellular evidence for the head-to-head dimerization is currently limited based on the available FoxP3 ChIP or CNR-seq data. Additionally, our new data showed that previously reported mutations that disrupt the head-to-head dimerization also affected the ladder-like multimerization, further limiting the ability to probe the physiological functions of the head-to-head dimerization. Nevertheless, given that head-to-head dimerization is unique to FoxP3, while the ladder-like multimerization is shared among all four FoxP TFs, we speculate that both forms exist in cells and carry out distinct functions depending on the bound DNA sequence. For example, DNA bridging would be a unique consequence of the ladder-like assembly, not shared with the head-to-head dimer, while the head-to-head dimerization may allow recruitment of certain cofactors using the unique surface created by the dimerization. This fits the previous microscopic analysis where FoxP3 were found in two distinct types of nuclear clusters associated with different cofactors^16^. Altogether, these findings suggest FoxP3 as a versatile TF that can interpret a wide range of sequences by assembling at least two distinct homo-multimeric structures.

Our findings also implicate functional roles of microsatellites in FoxP TF-mediated transcription regulation. While widely used as genetic tracing markers due to their high degrees of polymorphism, reports of biological functions of microsatellites^30,31^, besides its well-known pathogenic roles^32-35^, remain sparse^36,37^. Our finding of the T_n_G repeat recognition by FoxP3 and other members of the FoxP family, raises the question of whether microsatellites play greater and more direct roles in transcriptional regulation than previously thought. This also prompts speculation that microsatellite polymorphism may contribute to a broad spectrum of diseases through FOXP TF dysregulation, such as autoimmunity via FOXP3, neurodevelopmental disorders via FOXP1, speech and language impairments via FOXP2, and heart and hearing defects via FOXP4.

## EXPERIMENTAL MODEL AND SUBJECT DETAILS

### Mice

C57BL/6N mice, sourced from Taconic Biosciences and overseen by Harvard Medical Area (HMA) Standing Committee on Animals, were housed in an individually ventilated cage system at the specific-pathogen-free New Research Building facility of Harvard Medical School. The mice were maintained at a controlled environment with a temperature of 20-22°C, humidity ranging from 40-55%, and a 12-hour light-dark cycle. The spleens of 12∼14 weeks old female C57BL/6 mice were isolated for the study.

### Naive CD4+ T Cells

Cells were isolated by using Naive CD4^+^ T Cell Isolation Kit (Miltenyi Biotec, Cat#130-104-453) according to the manufacturer’s instructions and maintained in complete RPMI medium (10% FBS heat-inactivated, 2mM L-Glutamine, 1mM Sodium Pyruvate, 100μM NEAA, 5mM HEPES, 0.05mM 2-ME).

### HEK293T & A549 cells

HEK293T cells (were purchased from ATCC (CRL-11268)) and A549 cells (kind gift from Dr. Susan Weiss (U Penn)) were maintained in DMEM (High glucose, L-glutamine, Pyruvate) with 10% fetal bovine serum, 1% penicillin/streptomycin.

### EL4 cells

Cells (gift from Dr. Christophe Benoist lab (Harvard Medical School)) were cultured in DMEM (High glucose, L-glutamine, Pyruvate) supplemented with 10% fetal bovine serum, ranging from 1X10^5^ to 1X10^6^ cells/mL.

## METHOD DETAILS

### Plasmids

Mouse FoxP3 plasmids were made as previously described^22^. For Mammalian expression plasmids, HA-tagged mouse FoxP3 CDS was inserted into pcDNA3.1+ vector between KpnI and BamHI sites. All FoxP3 mutations including R356E, V396E, V398E, V408M and 409-411AAA were generated by site-directed mutagenesis using Phusion High Fidelity (New England Biolabs) DNA polymerases. For retroviral packaging plasmids, HA-tagged mouse FOXP3 CDS was inserted into MSCV-IRES-Thy1.1 vector.

For mammalian expression plasmids of FoxP3 orthologs from *H. sapiens, O. anatinus and D. rerio*, respective FoxP3 CDS with overhangs of pcDNA vector was synthesized by IDTDNA and then assembled with NEBuilder® HiFi DNA Assembly Cloning Kit (NEB, #5520G). FoxP3 paralogs (FoxP1, FoxP2 and FoxP4) mammalian expression plasmids were made in the same way. Other forkhead TFs, such as FoxA1, FoxM1, FoxQ1 and FoxS1, were gifts from Stefan Koch lab^40^ through Addgene.

### DNA oligos

Single-stranded DNA (ssDNA) oligos were synthesized by IDTDNA. Double-stranded DNA (dsDNA) oligos for EMSA assay, pulldown assay and DNA bridging assay were annealed from single-stranded, complementary oligos. After briefly spinning down each oligonucleotide pellet, ssDNAs were dissolved in the annealing buffer (10 mM Tris-HCl pH 7.5, 50 mM NaCl). Complementary ssDNAs were then mixed together in equal molar amounts, heated to 94°C for 2 minutes and gradually cooled down to room temperature. For dsDNA in cryo-EM analysis, HPLC purified single-stranded, complementary oligos were purchased from IDTDNA. After annealing, dsDNA was further purified by size-exclusion chromatography on Superdex 75 Increase 10/300 (GE Healthcare) columns in 20 mM Tris-HCl pH 7.5, 150 mM NaCl. Biotin labeled ssDNA oligos were synthesized by IDTDNA and then dissolved in annealing buffer (10 mM Tris-HCl pH 7.5, 50 mM NaCl). Complementary, biotin-labeled ssDNAs were then mixed together in equal molar amounts, heated to 94°C for 2 minutes and gradually cooled down to room temperature. Sequence of all the DNA oligos used are shown in Supplementary Table 2b.

### Protein expression and purification

All recombinant proteins in this paper were expressed in BL21(DE3) at 18°C for 16-20 hrs following induction with 0.2 mM IPTG. Cells were lysed by high-pressure homogenization using an Emulsiflex C3 (Avestin). All proteins are from the *Mus. musculus* sequence, unless mentioned otherwise. FoxP3^ΔN^ (residues 188-423) was expressed as a fusion protein with an N-terminal His_6_-NusA tag. After purification using Ni-NTA agarose, the protein was treated with HRV3C protease to cleave the His_6_-NusA-tag and were further purified by a series of chromatography purification using HiTrap Heparin (GE Healthcare), Hitrip SP (GE Healthcare) and Superdex 200 Increase 10/300 (GE Healthcare) columns. The final size-exclusion chromatography (SEC) was done in 20 mM Tris-HCl pH 7.5, 500 mM NaCl, 2 mM DTT. NFAT1 protein (residues 394-680) was also expressed as a fusion protein with an N-terminal His_6_-NusA tag. After purification using Ni-NTA agarose, the His_6_-NusA-tag was removed using the HRV3C protease and was further purified by SEC on Superdex 75 Increase 10/300 (GE Healthcare) column in 20 mM Tris-HCl pH 7.5, 500 mM NaCl, 5% Glycerol, 2 mM DTT. His_6_-MBP fused FoxP3^ΔN^ variants were purified using Ni-NTA affinity column and Superdex 200 Increase 10/300 (GE Healthcare) SEC column in 20 mM Tris-HCl pH 7.5, 500 mM NaCl, 2 mM DTT.

### MBP-FoxP3^ΔN^ pulldown-seq (FoxP3 PD-seq)

Mouse EL4 genomic DNA was isolated by QIAGEN Blood & Cell Culture DNA Kit (QIAGEN, #13343). The purified genomic DNA was then fragmented to 100∼200 bp using DNase I (ZYMO RESEARCH, #E1010) in the digestion buffer (50 mM NaCl, 20 mM Tris-HCl PH 7.5, 1.5 mM MgCl_2_) (for 200μL system, 50μg genomic DNA was treated with 8μL DNase I for 3∼4mins to obtain 100∼200 bp DNA fragments). The digested genomic DNA was then purified by QIAquick Nucleotide Removal Kit (QIAGEN, #28306) and used as an input for the pulldown-seq.

Purified MBP-tag or MBP-FoxP3^ΔN^ protein was incubated with the input DNA fragments in the incubation buffer (20 mM Tris-HCl pH 7.5, 100 mM NaCl, 1.5 mM MgCl_2_) for 20 mins at room temperature and then subjected to MBP pulldown using Amylose Resin (New England Biolabs). The bound DNA was recovered using proteinase K (New England Biolabs) and purified using QIAquick Nucleotide Removal kit (QIAGEN). The sequencing libraries were made using NEBNext® Ultra™ II DNA Library Prep Kit (Illumina) according to the manufacturer’s instructions and submitted to Novogene for NGS with PE150.

### Nucleosome pulldown-seq

Mouse EL4 cells were lysed using a hypotonic buffer (20mM Bis-Tris pH 7.5, 0.05% NP-40, 1.5mM MgCl_2_, 10mM KCL, 5mM EDTA, 1Xmammalian protease inhibitor) and the nuclear fraction was isolated by spinning down at 4° and 2,500 RPM for 10 mins. The isolated nuclear fraction was then digested with Micrococcal Nuclease (Thermo Scientific, #88216) for 1 hour at 4℃ to fragment the chromatin into individual nucleosomes. Lysate was then spun down at 4° and 13,000 RPM for 10 mins. Cleared lysate containing the nucleosomes was incubated with purified MBP-tag or MBP-FoxP3^ΔN^ protein (1uM) for 1 hour at 4℃ and then subjected to MBP pulldown using Amylose Resin (New England Biolabs). After proteinase K (New England Biolabs) treatment, the final nucleosomal DNAs were recovered by QIAquick Nucleotide Removal kit (QIAGEN) and used for library preparation. The libraries were made by NEBNext® Ultra™ II DNA Library Prep Kit (Illumina) according to the manufacturer’s instructions and submitted to Novogene for NGS with PE150.

### MBP-FoxP3^ΔN^ pulldown assay

0.4 μM purified MBP-mFoxP3^ΔN^ protein was incubated with 0.1 μM DNA in the incubation buffer for 20 mins. The FoxP3-DNA mixture was then incubated with 25μL Amylose Resin (New England Biolabs) for 30 mins with rotation at room temperature. The bound DNA was recovered using proteinase K (New England Biolabs), purified using QIAquick Nucleotide Removal kit (QIAGEN) and analyzed on 10% Novex TBE Gels (Invitrogen). DNA was visualized by Sybr Gold staining. The expression of MBP-FoxP3^ΔN^ was validated by Western blot using Mouse MBP Tag antibody(8G1) (Cell Signaling Technology, Cat#2396, 1:2000 dilution).

### HA-FoxP3 pulldown assay

HEK293T cells were transfected with pcDNA encoding HA-tagged FoxP3 (wild-type or mutants). After 48 hours, cells were lysed using RIPA buffer (10mM Tris-HCl pH 8.0, 1mM EDTA, 1% Triton X-100, 0.1% Sodium Deoxycholate, 0.1% SDS, 140 mM NaCl and 1X proteinase inhibitor) and treated with Benzonase (Millipore) for 30 mins. The lysate was then incubated with Anti-HA Magnetic Beads (Thermo Fisher) for 1 hour. Beads were washed three times using RIPA buffer and incubated with DNA oligos for 20 mins at room temperature. Bound DNA was recovered using proteinase K (New England Biolabs), purified using QIAquick Nucleotide Removal kit (QIAGEN) and analyzed on 10% Novex TBE Gels (Invitrogen). DNA was visualized by Sybr Gold staining.

### Nucleosome reconstitution and EMSA analysis

Nucleosome core particles (NCP) were reconstituted with recombinant Histone Octamer H3.1 (Active motif) and DNAs as described previously^41^. Briefly, 1 μM of TTTG repeats (144bp), AAAG repeats (144bp), TGTG repeats (144bp) and DNA harboring 601 sequence (181bp) were incubated with 1 μM of the histone octamer and were dialyzed against 10mM Tris-HCl PH 7.5, 1mM EDTA, 2mM DTT for 24 hrs. Nucleosomes (0.05 μM) were incubated with the indicated amount of FoxP3^ΔN^ in the buffer (10mM Tris-HCl pH 7.5, 50mM NaCl, 1mM EDTA, and 2mM DTT) for 30 min at 4 °C and analyzed on 6% TBE gels (Life Technologies) at 4 °C. After staining with Sybr Gold stain (Life Technologies), Sybr Gold fluorescence was recorded using iBright FL1000 (Invitrogen) and analyzed with iBright Analysis Software.

### Biotin-DNA pulldown assay

HA-FoxP3 was transiently expressed in HEK293T cells as above. Cells were lysed using RIPA buffer. The lysate was incubated with biotin-dsDNA (1μM) for 1 hour, and then with streptavidin agarose beads (Thermo Fisher, 25μL) for additional 30mins. Beads were spun down and washed three times with RIPA buffer. Bead-bound protein was extracted using the SDS loading buffer and analyzed by SDS-PAGE and western blotting using anti-HA (Primary) antibody (Cell Signaling, Cat#3724S, 1:3000 dilution) and anti-rabbit IgG-HRP (Secondary) antibody (Cell Signaling, Cat#7074, 1:5000 dilution).

### Electrophoretic Mobility Shift Assay (EMSA)

DNA (0.05μM) was mixed with the indicated amount of FoxP3 in buffer A (20mM HEPES pH 7.5, 150mM NaCl, 1.5mM MgCl_2_ and 2mM DTT), incubated for 30 min at 4 °C and analyzed on 3-12% gradient Bis-Tris native gels (Life Technologies) at 4 °C. After staining with Sybr Gold stain (Life Technologies), Sybr Gold fluorescence was recorded using iBright FL1000 (Invitrogen) and analyzed with iBright Analysis Software.

### Crosslinking analysis

Protein-protein crosslinking using BMOE (Thermo Scientific) was carried out according to the product manual. Briefly, 0.4μM FoxP3 ^ΔN^ was incubated with 0.05μM DNAs at 25°C for 10 mins in 1X PBS, then BMOE was added to the final concentration of 100 μM. After 1 hour incubation at 25°C, DTT (10mM) was added to quench the crosslinking reaction. Samples were then analyzed by SDS-PAGE and Krypton staining (Thermo Scientific).

### DNA bridging assay

Biotin-DNA (bait, 0.1μM) was incubated with Streptavidin Agarose (25μL, Thermo Fisher) in buffer B (20mM Tris-HCl pH 7.5, 100mM NaCl, 1.5mM MgCl_2_, 5mM DTT) for 30 mins by rotating the mixture at room temperature. Agarose beads were washed three times with buffer B and incubated with non-biotinylated DNA (prey, 0.1μM) and purified FoxP3 protein (or 293T lysate expressing FoxP3). After incubation for 30 mins with rotation, bead-bound DNA was recovered using proteinase K (New England Biolabs), purified by QIAquick Nucleotide Removal kit (QIAGEN) and analyzed on 10% Novex TBE Gels (Invitrogen). DNA was visualized by Sybr Gold staining.

### Cryo-EM sample preparation and data collection

FoxP3^ΔN^ was incubated with (T_3_G)_18_ DNA at a molar ratio of 8:1 in buffer B at RT for 10 mins. The complex was then crosslinked using 0.5 % glutaraldehyde for 10 mins at RT prior to quenching with 1/10 volume of 1M Tris-HCl pH 7.5 (for a final Tris concentration of 0.1M). The FoxP3^ΔN^-DNA complex was then purified by Superose 6 Increase 10/300 GL (GE Healthcare) column in 20mM Tris-HCl pH 7.5, 100mM NaCl, 2mM DTT. The sample was concentrated to 1 mg/ml (final for protein) and applied to freshly glow-discharged C-flat 300 mesh copper grids (CF-1.2/1.3, Electron Microscopy Sciences) at 4℃. The grids were plunged into liquid ethane after blotting for 5 s using Vitrobot Mark IV (FEI) at the humidity setting of 100%. The grids were screened at the Harvard Cryo-EM Center and UMass Cryo-EM core facility using Talos Arctica microscope (FEI). The grids that showed a good sample distribution and ice thickness were used for data collection on Titan Krios (Janelia Cryo-EM facility) operated at 300 kV and equipped with Gatan K3 camera. A total of 11624 micrographs were taken at a magnification of 81,000x with a pixel size of 0.844 Å. Each movie comprised of 60 frames at total dose of 60 e^-^/Å^2^. The data were collected in a desired defocus range of -0.7 to -2.1 mm.

### Cryo-EM data processing and structure refinement

Data were processed using cryoSPARC V4.2.0^42^ and RELION V4.0.1^43,44^. The dose-fractionated movies were motion corrected using MotionCor2^45^. The contrast transfer function (CTF) was estimated with CTFFIND 4.1^46^. Particles were picked using the Auto pick function in RELION^47^. 4,201,166 raw particles were transferred to cryoSPARC for 2D classification. 1,009,168 particles from selected 2D classes were used for Ab-initio reconstruction, where they were divided into 6 Ab-initio classes. 317,175 particles from class 1 were then refined to a final resolution of 3.7 Å with non-uniform refinement. To improve local resolution, we performed local refinement using a mask covering the central FoxP3 tetramer, and obtained a 3.3 Å-resolution map. For structure refinement, a previous crystal structure of a FoxP3^ΔN^ monomer bound to DNA (PDB: 7TDX) was docked into the EM density map from global refinement using UCSF Chimera^48^. The total of ten copies of FoxP3^ΔN^ monomers were located for the global refinement map. For the mask-focused local refinement map, four copies of FoxP3^ΔN^ monomers in complex with DNA were docked. Subsequently, the decamer and tetramer models were built manually against the respective density map using COOT^49^, and refined using phenix.real_space_refine^50^. The structure validation was performed using MolProbity^51^ from the PHENIX package. The curve representing model versus full map was calculated, based on the final model and the full map. The statistics of the 3D reconstruction and model refinement are summarized in Extended Data Table 1. All molecular graphics figures were prepared with Pymol (Schrödinger, LLC) and UCSF Chimera^48^. All softwares used for cryo-EM data processing and model building were installed and managed by SBGrid^52^.

### Negative stain EM

FoxP3^ΔN^ (0.4μM) was incubated with DNA (0.05μM) in buffer B at RT for 10 mins. The samples were diluted 10-fold with buffer A, immediately adsorbed to freshly glow-discharged carbon-coated grids (Ted Pella) and stained with 0.75% uranyl formate as described before^53^.sImages were collected using a JEM-1400 transmission electron microscope (JEOL) at 50,000X magnification.

### *De novo* motif analysis of FoxP3-occupied sites *in vitro* and *in vivo*

FoxP PD-seq data were mapped to mm10 using Bowtie2^54^ and sorted by samtools^55^. Peaks were called using MACS2^56^ with either input or MBP pull-down as controls. Default setting was used for peak calling. *De novo* motif analysis was done using MEME-ChIP^57^ and STREAM^58^ with minimum and maximum motif lengths set at 6 and 30 nt, respectively. FoxP3 CUT&RUN-seq and ChIP-seq data^14^ were mapped to mm10 using Bowtie2^54^. Peaks were called using MACS2^56^. Bedtools was used to obtain CNR -seq consensus (n=1,372) and union (n=9,062) peaksbetween Rudensky and Dixon CNR peaks. Motif analysis was done as above. To independently validate the results, similar motif analysis was repeated using different ChIP-seq data^23,24^, which were mapped to mm10 using Bowtie2. Peaks were called using HOMER with an input control^22^ and ranked them based on the signal intensity using samtools^55^. Top 5,000 overlapping FoxP3 ChIP-seq peaks were calculated by bedtools using a 50% reciprocal overlap criterion. FoxP3-negative open chromatin regions (OCRs) were derived from all observed Treg OCRs^27^. Intersections and non-overlapping genomic features were extracted using bedtools^59^ *intersect* functionality and were subjected to the motif analysis as above. The versions and parameters for softwares used above have been uploaded to GitHub (https://github.com/DylannnWX/Hurlab/tree/main/Foxp3_manuscript).

### Genome-wide analysis of TnG repeat-like elements

FIMO^60^ was used to identify TnG repeat-like elements. The TnG repeat-like motif identified from the MEME-ChIP analysis of the overlap of Rudensky and Dixon CNR peaks (Supplementary Table 1b) was used as a query motif, and search was done against the human (GrCh38), mouse (GrCm38), and Zebrafish (GrCz11) genomes. The default p-value cutoff (p=0.05) was used. FIMO outputs of all regions that match the query motif were converted to bed file format, and the overlapping TnG regions from FIMO outputs were combined into a single region by bedtools merge function.

### Comparison between FoxP3 CNR union peaks with and without T_n_G repeat-like elements

FIMO^60^ was used as above to identify TnG repeat-containing peaks from the Rudensky/Dixon CNR union peaks (n=9,062). Out of the 9,062 peaks, 3,301 peaks showed at least one T_n_G region lower than the default p-value cutoff (p=0.05), and were classified as T_n_G-containing peaks. The non-TnG-containing peaks were then calculated by bedtools peak subtraction with intersect -v. Genomic feature analysis was done using ChIPseeker^61^. To compare H3K4me3, H3K27ac and ATAC signal intensity, H3K4me3 and H3K27ac ChIP-seq and ATAC-seq data^23^ were mapped to mm10 using Bowtie2^54^ and the intensity was calculated within 2 kb upstream and downstream of the FoxP3 CNR peak summits using Deeptools^62^ bamCoverage and Deeptools computeMatrix. The versions and parameters for softwares used above have been uploaded to GitHub (https://github.com/DylannnWX/Hurlab/tree/main/Foxp3_manuscript).

### Motif analysis of other Forkhead TFs

Peak bed files for Foxp1, Foxp2, Foxp4, Foxj2, Foxj3, FoxA1, FoxM1, FoxS1 and FoxQ1, were from downloaded from ChIP-Atlas (http://chip-atlas.org/) and converted to fasta files using bedtools^59^ getfasta. The individual fasta file was then subjected to *de novo* motif analysis using MEME-ChIP^57^ with the minimum and maximum motif lengths set at 6 and 30 nt, respectively. The results are summarized in Supplementary Table 1c.

### CD4^+^ T Cell isolation and retroviral transduction

Naïve CD4^+^ T cells were isolated by negative selection from mouse spleens using the isolation kit (Miltenyi Biotec) according to the manufacturer’s instruction. The purity was estimated to be >90% as measured by PE anti-CD4 (Biolegend, Cat#100408, 1:1000 dilution) staining and FACS analysis. Naïve CD4^+^ T cells were then activated with anti-CD3 (Biolegend, Cat#100340, 1:500 dilution to 5μg/mL), anti-CD28 (Biolegend, Cat#102116, 1:500 dilution to 5μg/mL) and 50 U/mL of IL2 (Peprotech) in complete RPMI medium (10% FBS heat-inactivated, 2 mM L-Glutamine, 1 mM Sodium Pyruvate, 100 μM NEAA, 5 mM HEPES, 0.05 mM 2-ME). The activation state of T cells was confirmed with increased cell size and CD44 (BioLegend) expression by FACS. After 48 hours, cells were spin-infected with retrovirus containing supernatant from HEK293T cells transfected with retroviral expression plasmids (Empty MSCV-IRES-Thy1.1 vector, wildtype-FoxP3 and mutations encoding vectors) and cultured for 2∼3 days in complete RPMI medium with 100 U/mL of IL2.

### FoxP3 transcriptional activity assay in CD4^+^T cells

FoxP3 transcriptional activity was measured by levels of two known target genes, CD25 and CTLA4, and the FoxP3 expression marker Thy1.1. FoxP3-transduced CD4^+^ T cells were stained with antibodies targeting the cell surface antigens CD25 (Biolegend, Cat#102022, 1:1000 dilution) and Thy1.1 (Biolegend, Cat#202520, 1:1000 dilution) on day 2 post-retroviral infection. The level of CTLA4 was measure by intracellular staining using anti-CTLA4 (Biolegend, Cat#106311, 1:1000 dilution) and the Transcription Factor Staining Buffer Set (eBioscience) on day 3 post retroviral infection. Flow cytometry data were analyzed with FlowJo software and presented as plots of mean fluorescence intensity (MFI) of CD25 and CTLA4 in cells grouped into bins of Thy1.1 intensity, which is the expression marker for FoxP3. Each result is representative of 3 independent experiments.

### FoxP3 ChIP-seq analysis

FoxP3 ChIP-seq was conducted using CD4^+^ T cells following a published procedure^63^. Activated CD4^+^ T cells that had been transduced with wild-type or mutant FoxP3 were sorted based on Thy1.1 reporter expression. For each sample (5X10^6^ cells), cross-linking was achieved with 1% formaldehyde for 10 minutes. Subsequently, the cells were lysed on ice using RIPA buffer (10mM Tris-HCl pH 8.0, 1mM EDTA, 1% Triton X-100, 0.1% Sodium Deoxycholate, 0.1% SDS, 140 mM NaCl, and 1X proteinase inhibitor). Chromatin fragmentation was achieved using an AFA Focused-ultrasonicator (Covaris M220) for 30 minutes (5% duty cycle, 140W max power, 200 cycle/burst), resulting in DNA fragments ranging from 100 to 200 bp. The sheared material underwent a 10-minute centrifugation at 13,000 rpm at 4°C to clear the solution. The cleared material was then subjected to immunoprecipitation overnight with HA-tag antibody (Cell Signaling, #3724) at 4°C, and Protein G beads (Active motif, #53014) were added for an additional 2 hours. The beads were sequentially washed with various buffers: RIPA Wash Buffer (0.1% SDS, 0.1% Sodium Deoxycholate, 1% Triton X-100, 1mM EDTA, 10mM Tris-HCl pH 8.0, 150mM NaCl), RIPA 500 Wash Buffer (0.1% SDS, 0.1% Sodium Deoxycholate, 1% Triton X-100, 1mM EDTA, 10mM Tris-HCl pH 8.0, 500mM NaCl), LiCl wash buffer (10mM Tris-HCl, pH 8.0, 250mM LiCl, 0.5% Triton X-100, 0.5% Sodium Deoxycholate), and Tris buffer (10mM Tris-HCl, pH 8.5). The chromatin was eluted from the beads using Elution buffer (1X TE, pH 8.0, 0.1% SDS, 150mM NaCl, 5 mM DTT). After elution, the DNA was treated with 1 µg DNase-free RNase (Roche) for 30 minutes at 37°C, followed by treatment with Proteinase K (Roche) for at least 4 hours at 63°C to reverse the crosslinks. The reverse-crosslinked DNA was then purified using SPRI beads (Beckman, #B23318). Subsequent steps, including end repair, A-base addition, adaptor-ligation, and PCR amplification, were carried out to prepare the ChIP-seq library for each sample. The libraries were made by NEBNext® Ultra™ II DNA Library Prep Kit (Illumina) according to the manufacturer’s instructions and submitted to Novogene for NGS with PE150.

### mRNA-seq analysis

mRNA-seq was conducted using CD4^+^ T cells. Activated CD4^+^ T cells that had been transduced with wild-type or mutant FoxP3 were sorted based on Thy1.1 reporter expression. For each sample, 1X10^6^ cells were sorted and subjected to total RNA extraction using TRIzol reagent and Direct-zol RNA Miniprep Kits. Quality control and the construction of mRNA-seq libraries were undertaken by Novogene Co. Ltd.. The NEB Next Ultra II kit and the non-directional mRNA approach with the polyA pipeline were used. The libraries were subsequently sequenced using the Illumina NovaSeq 6000 instrument, generating paired-end reads with a length of 2X150 bp, resulting in ∼30 M reads per sample. Raw sequence files were subjected to pre-processing using Trimmomatic v.0.36 to remove Illumina adaptor sequences and low-quality bases. Trimmed reads were then aligned to the mouse genome (UCSC mm10) using bowtie2/2.3.4.3. For gene read counting, HTseq-count (v. 0.12.4) was employed. Normalization of gene counts and differential analysis were carried out using DESeq2 (v. 5). The creation of heatmaps was accomplished using Pheatmap.

### Chromatin contact analysis

HiC- and PLAC-seq datasets were downloaded from GSE217147^12^, and the list of Treg enhancer-promoter loops (EPLs) was obtained from^13^. All .hic files were converted to .cool files to hic2cool, and all .cool files were decompressed into .txt files by cooler dump –join function. These decompressed files were loaded as python pandas dataframes. All possible bins in .cool files were converted to bed file formats, and intersected with TnG-containing or TnG-absent CNR union peaks using bedtools intersect -wa function to acquire the bins that contain TnG bins and non-TnG (NTnG) bins. These bins were used as anchors to filter raw .cool files for contact pairs between TnG–TnG (2TnG), TnG–NTnG (TnGNTnG), and NTnG–NTnG (2NTnG). These contact pairs were then filtered by [more than 5 in WT Treg HiC-seq] and [more than indicated threshold in FoxP3 PLAC-seq]. Contact counts in Fig. 2f were listed in Supplementary Table 3.

The p-value of 2TnG pair enrichment was first calculated by getting the expected 2TnG pair counts in a given list of pairs assuming random distribution (number of contact pairs * proportion of all potential TnG bins^2). Then, this number was compared with the observed 2TnG pair counts using binomial distribution. The proportion of all potential TnG bins is 0.37, which matches the proportion of TnG CNR peaks out of all CNR peaks (3,301 out of 9,062). The p-value was the cumulated probability that the observed 2TnG pair counts happen by chance, and the alternative hypothesis, if the p-value is low, indicates the probability that in 2TnG pair is enriched in the given list of contact.

To compare HiC/PLAC-seq anchors (in mm9) to EPL anchors (in mm10), the reference genomes of mm9 were lifted to mm10 using UCSC genome browser to acquire the correlating bin coordinates in mm10, and their overlaps were analyzed using bedtools intersect function.

### T cell suppression assay

Isolated naïve CD4^+^ T cells were activated with anti-CD3 (Biolegend), anti-CD28 (Biolegend) and 50 U/mL of IL2 (Peprotech) in complete RPMI medium. After 48 hours, activated CD4^+^ T cells were retrovirally transduced to express FoxP3 and were used as “suppressors”. In parallel, freshly isolated naïve CD4^+^ T cells were labelled with CellTrace™ CFSE (Invitrogen) and used as “responders”. CD3^-^ T cells representing APC cells were also isolated using the isolation kit (Miltenyi Biotec) according to the manufacturer’s instructions. For suppression assay, the CFSE-labeled responder cells (5X10^4^ cells) were stimulated with APC cells (10^4^ cells) and anti-CD3 (1 μg/mL) in 96-well round-bottom plates for 3 days, in the presence or absence of FoxP3-transduced suppressor cells (at a responder-to-suppressor ratio of 2:1). Proliferation ratio of the responders were calculated as a function of CFSE dye dilution by FACS analysis.

### Statistics and Reproducibility

Data in Figures 1f-j, 2e, 3b-e, 4b, 4d, 4e, 4f, 4g, 5b, 5c and extended data figures 1g-l, 2a, 2h, 3b, 3c, 3e, 3f, 5c-e, 6a-e, 6g are representative of at least three independent experiments and each experiment was repeated independently with similar results.

## Data availability

Naked genomic DNA pulldown-seq, nucleosome pulldown-seq, FoxP3 mRNA-seq and FoxP3 ChIP-seq data have been deposited to the GEO database with the accession code of GSE243606. The structures and cryo-EM maps have been deposited to the PDB and the EMDB under the accession codes of 8SRP and EMD-40737 for the decameric FoxP3 in complex with DNA, and 8SRO and EMD-40736 for the central FoxP3 tetramer in complex with DNA (focused refinement). Other research materials reported here are available on request.

## Code availability

All custom codes used in this project are deposited to GitHub (link: https://github.com/DylannnWX/Hurlab/tree/main/Foxp3_manuscript). These include the processing of Deeptools matrix outputs, FIMO region to peak bed files, and HiC/Cool data processing. All are standalone Jupyter Notebook instances. In each instance, detailed user instructions, example inputs and expected outputs were also included in this GitHub Repo.

## Acknowledgements

We thank all members of the Hur lab, A. Rudensky (MSKCC) and Y. Zhong (Shanghai Jiao Tong University) for their helpful discussion and feedback. This study was supported by Modell fellowship to W.Z., NIH grants (R01AI180137, R01AI154653 and R01AI111784 to S.H; AI150686 and AI165697 to C.B.) and Howard Hughes Medical Institute (S.H.). Cryo-EM data were collected at Cryo-EM facilities at Janelia, Harvard Medical School and University of Massachusetts Worcester.

## Author contributions

W.Z., F.L., S.H. conceived and designed the project. W.Z., F.L. performed all the experiments. F.L. determined the structure. W.Z., X.W, R.R. performed bioinformatic analysis. J.P. assisted experiments. C.B. and S.H. supervised bioinformatic analysis. S.H. supervised overall project.

## Competing Interests

The authors declare no competing interests.

## Extended figure legends

**Extended Data Figure 1. Analysis of T_3_G repeats in the genome and FoxP3 multimerization on T_3_G repeats**

a. TnG repeat-like sequences in the genomes of *H. sapiens*, *M. musculus* and *D. rerio*. Sequences that match the TnG repeat-like motif (29 nt motif from FoxP3 CNR overlap peaks, see Supplementary Table 1b) were identified using FIMO (*p* = 0.05, see Methods). Genomic percentage of TnG repeat-like sequences (in parenthesis) was the number of TnG repeat regions multiplied by the average size of the repeats (31, 33 and 38 bp for *H. sapiens*, *M. musculus* and *D. rerio*, respectively), divided by the genome size (3.2, 2.7 and 1.4 billion bp, respectively). Below: length distribution of the TnG repeat-like sequences. Genes-to-genome size ratio was calculated by dividing the number of genes used in the feature annotation (31,074, 24,528 and 13,576 in *H. sapiens*, *M. musculus* and *D. rerio*) by the genome size.
b. Distribution of TnG repeat-like sequences in the genomes of *H. sapiens*, *M. musculus* and *D. rerio* relative to Transcription Start Sites (TSSs).
c. Distribution of CNR union peaks (union of Rudensky CNR peaks and Dixon CNR peaks, n=9,062) relative to TSSs. CNR union peaks with and without TnG repeat-like sequences (n=3,301 and 5,761, respectively) were identified using FIMO (*p* = 0.05) as in (a).
d-f. Comparison of (d) H3K4me3-ChIP, (e) H3K27ac-ChIP and (f) ATAC signal ^23^ around the TnG repeat-like sequences that overlap with FoxP3 CNR union peaks vs. those genome-wide in thymic Tregs (n = 4,837 peaks and 41,889 peaks respectively). See Extended Data Figs. 4a-4c for pre-thymic Tregs, which showed that high levels of H3K4me3, H3K27ac and ATAC signals were maintained prior to FoxP3 expression. TnG repeats in the blacklist were removed. Right: ChIP/ATAC signal was averaged over +/- 500 bp around the TnG repeat-like sequences. Two-tailed unpaired t-tests. ****, *p* < 0.0001.
g. DNA sequence specificity of FoxP3 as measured by FoxP3 pull-down. HA-tagged, full-length FoxP3 was transiently expressed in HEK293T cells and purified by anti-HA IP. Equivalent amounts of indicated DNAs (30-31 bp) were added to FoxP3-bound beads and further purified by anti-HA IP prior to gel analysis.
h. DNA sequence specificity of FoxP3 as measured by DNA pull-down. Equivalent amounts of biotinylated DNAs were mixed with FoxP3-expressing 293T lysate and were subjected to streptavidin pull-down. Co-purified FoxP3 was analyzed by anti-HA WB.
i. FoxP3 binding to nucleosomal DNA as measured by native gel-shift assay. Indicated DNA was incubated with the histone octamer at 1:1 molar ratio (black circle), followed by incubation with FoxP3 (0.2 or 0.4 μM for light and dark green circles, respectively). Empty dotted circles indicate no histone or FoxP3. Sybrgold stain was used for visualization. With an increasing concentration of FoxP3, the intensity of the nucleosomal TTTG repeat decreased, while the signal in the gel well (red arrow) increased. Such changes were not observed with other DNAs.
j. BMOE crosslinking of FoxP3^ΔN^ with and without DNA. FoxP3^ΔN^ can only form multimers on (T3G)6 DNA.
k. Multimerization analysis of FoxP3, as measured by co-purification of FoxP3 with different tags. GST- and MBP-tagged FoxP3 were incubated together in the presence and absence of indicated DNA and were subjected to MBP pull-down, followed by WB analysis of GST-FoxP3 in eluate. Note that GST replaced the CC domain in FoxP3, disallowing hetero-dimerization between MBP-FoxP3 and GST-FoxP3. Thus, co-purification of these two proteins in the presence of T3G repeats suggests DNA sequence-dependent multimerization of the FoxP3 homodimer.
l. Representative negative-stain EM images of FoxP3^ΔN^ in complex with (AAAG)36 (left) and (TGTG)36 (right).

**Extended Data Figure 2. Cryo-EM structure of the FoxP3^ΔN^–(T_3_G)_18_ complex.**

a. Representative negative-stain EM (left) and cryo-EM images (right) of FoxP3^ΔN^ multimers on (T3G)18 DNA.
b. 2D classes chosen for 3D reconstruction.
c. Cryo-EM image processing workflow. See details in Methods.
d. Local resolution for the maps of global refinement (left) and local refinement with a mask covering the central four subunits of FoxP3 (right). Local resolution was calculated by CryoSPARC. Resolution range was indicated according to the color bar.
e. Fourier shell correlation (FSC) curve for global refinement (left) and local refinement (right). Map-to-Map FSC curve was calculated between the two independently refined half-maps after masking (blue line), and the overall resolution was determined by gold standard FSC=0.143 criterion. Map-to-Model FSC was calculated between the refined atomic models and maps (red line).
f. Cryo-EM map and ribbon model of FoxP3^ΔN^ decamer in complex with two (T3G)18 DNAs (PDB: 8SRP, EMDB: 40737). DNA molecules are colored grey. Individual FoxP3 monomers are colored differently.
g. Superposition of the domain-swap dimeric structure of FoxP3 (cyan, PDB:4WK8) onto any subunit of the FoxP3 multimeric structure (represented here by the orange subunit) by aligning the common portions of FoxP3 reveals that the domain-swap dimer is incompatible with the density map.
h. Native gel shift analysis of MBP-tagged FoxP3^ΔN^ (WT or R337Q, 0.4 μM) with (T3G)12 DNA (0.05 μM). Note that R337Q induces domain-swap dimerization.

**Extended Data Figure 3. DNA sequence and conformational analysis.**

a. DNA sequence assignment by Q-score analysis using mapQ. For each of the four DNA strands, eight possible sequence alignments (right) were tested against the local refinement map. The sequence alignment showing the highest overall score was highlighted with a thick red line. The best sequence alignments for all four DNA strands were consistent with each other and were used for cryo-EM reconstruction.
b. Experimental validation of FoxP3–DNA registers using NFAT–FoxP3 cooperativity analysis. This assay utilizes the FoxP3 interaction partner NFAT, which assists FoxP3 binding to DNA only when their binding sites are 3 bp apart in one particular orientation (as in the schematic). To investigate FoxP3 footprints on T3G repeat DNA, we varied the position and orientation of NFAT consensus sequence (GGAAA, green) relative to T3G repeats (Var1-6), and performed FoxP3 pull-down. An internal control DNA (cntrl, harboring the NFAT motif followed by FKHM with 3 nt gap) was used to normalize the test DNA (Var1-6) pull-down efficiency. Only DNA with a single nucleotide gap between GGAAA and TTTG (Var2) showed a positive effect of NFAT on the FoxP3– DNA interaction. This suggests that the most upstream FoxP3 subunit recognizes TGTTTGT. Two-tailed paired t-tests, comparing with and without NFAT. *p*<0.005 for **, *p*<0.05 for * and *p*>0.05 for ns.
c. FoxP3 interaction with (T3G)10 variants. Variations in DNA sequence outside the FoxP3 footprints were tolerated (Var7 and Var8), but those within the footprints (Var9 and Var10) were not.
d. Comparison of the inter-subunit interactions in FoxP3 decamer on TnG repeats vs head-to-head (H-H) dimer on IR-FKHM (PDB:7TDX). Superposition of the H-H dimer (dark grey) onto any of the ten subunits in the decamer structure showed distinct modes of inter-subunit interactions. Shown are two examples where subunit 1 of the H-H dimer was aligned to orange (top) or magenta (bottom) subunits of the decamer, showing that subunit 2 of the H-H dimer did not align with any of the decamer subunits.
e. DNA bridging assay using FoxP3 expressed in 293T cells. Biotinylated and non-biotinylated DNA (82 and 60 bp, respectively) were mixed at 1:1 ratio and further incubated with 293T lysates expressing HA-tagged FoxP3, followed by streptavidin pull-down and gel analysis of non-biotinylated DNA by SybrGold staining.
f. DNA bridging assay using an increasing concentration of purified FoxP3^ΔN^. 0.1 μM each of biotinylated and non-biotinylated (T3G)12 DNAs were used. FoxP3 concentrations are indicated at the bottom.

**Extended Data Figure 4. FoxP3 can bridge T_n_G repeat-containing sites in vivo.**

a-c. Comparison of (a) H3K4me3-ChIP, (b) ATAC and (c) H3K27ac-ChIP signal ^23^ around the CNR union peaks with and without TnG repeat-like sequences in pre-thymic Tregs (pre-tTregs) and thymic Tregs (tTregs) (n=3,301 peaks and 5,761 peaks, respectively). Right: ChIP/ATAC signal was averaged over +/- 500 bp around the CNR peak summits. Two-tailed unpaired t-tests. ****, *p* < 0.0001.
d. FoxP3-dependence of the chromatin contacts at FoxP3-bound TnG anchors in Fig. 2f. FoxP3-bound TnG anchors were defined as anchors that overlap with FoxP3 CNR peaks with TnG repeat-like sequences. Contacts with frequency>5 in WT Treg HiC and connected by two TnG anchors were analyzed with an increasing FoxP3 PLAC-seq count threshold. For each contact, log2 foldchange of HiC counts from WT to FoxP3 knock-out Treg were plotted. Contacts with FDR<10 were colored red. The majority of the TnG–TnG contacts were less frequent in FoxP3 knock-out than in WT Tregs, although smaller fractions (10-15%) showed statistically significant FoxP3 dependence (FDR<10), as previously reported^12^. n=4365, 2559, 813, 204 and 60 anchors respectively. Mean ± SD were shown in black and blue lines. See also Supplementary Table 3.
e. Fraction of the FoxP3-bound TnG anchors from Fig. 2f that overlap with previously published Treg EPL anchors ^13^.

**Extended Data Figure 5. Characterization of intra-rung interface mutant FoxP3**

a. mRNA-seq heatmap analysis. CD4^+^ T cells were transduced and sorted to express FoxP3 as in Fig. 3d and were subjected to mRNA-seq. Top 100 genes showing the most significant difference between WT FoxP3 and EV were chosen for the evaluation of individual mutants. All four mutants were impaired in transcriptional functions, albeit to varying extents. The level of FoxP3 was equivalent for WT and all mutants. A few genes previously reported to be FoxP3-dependent were indicated in larger fonts. Note that V398E was not tested due to its negative effect on NFAT binding in (e).
b. ChIP-seq of HA-tagged FoxP3. Cells were transduced as in Fig. 3d and were subjected to anti-HA ChIP-seq. WT FoxP3 bound peaks were identified using MACS2 (n=8,607, *p*<0.01), and heatmaps of the ChIP signal were generated for each mutant at the WT peak locations. Below: averaged intensity of ChIP signal within 0.5 kb of the WT peak summit. Peaks with and without TnG repeats (n=1,900 peaks and 6,707 peaks, respectively) were compared. Two-tailed paired t-tests, comparing mutants to WT. *p* < 0.0001 for ****, *p*<0.001 for *** and *p*<0.005 for **.
c. Nuclear localization of WT FoxP3 and intra-rung interface mutants. HA-tagged FoxP3 was transiently expressed in A549 cells and was subjected to anti-HA immunofluorescent (yellow) analysis. Nuclei were shown with DAPI (blue) staining.
d. Expression levels of WT FoxP3 and intra-rung mutants in A549 cells.
e. Effect of the intra-rung mutations on the NFAT–FoxP3 interaction, as measured by native gel shift assay. FoxP3 (0.1 μM) was incubated with DNA harboring IR-FKHM and the NFAT site (with a 3-bp gap as in Extended Data Fig. 3b, 0.05 μM). NFAT (0.1 μM) was added to the mixture to monitor formation of the ternary complex NFAT– FoxP3–DNA. Note that V398E showed slight but reproducible reduction in NFAT binding.

**Extended Data Figure 6. Multimerization on T_n_G repeats is conserved in FoxP3 orthologs and paralogs.**

a. TnG repeats DNA-binding activity of FoxP3 with mutations in RBR. All seven RBR mutations previously shown to disrupt the head-to-head dimerization ^22^ disrupted (T3G)12 binding.
b. Effect of inter-rung^12bp^ spacing variations on FoxP3-mediated DNA bridging. Non-biotinylated DNAs in Fig. 4d were mixed with biotinylated (T3G)10 and FoxP3^ΔN^ (0.2 μM) prior to streptavidin pull-down and gel analysis. Relative level of non-biotinylated DNA co-purified with biotinylated DNA was quantitated from three independent pull- downs. Two-tailed paired t-tests, in comparison to (T3G)10. *p*<0.001 for ***, *p*<0.05 for * and *p*>0.05 for ns.
c. DNA-bridging activity of FoxP3 with different combinations of TnG repeats. Biotinylated-DNA (red) and non-biotinylated DNA (blue) were mixed at 1:1 ratio and were incubated with FoxP3^ΔN^ (0.2 μM) prior to streptavidin pull-down and gel analysis of non-biotinylated DNA. T2G, T4G and T5G repeats bridged better with T3G repeats than with themselves.
d. DNA-binding activity of FoxP3 orthologs with indicated DNA. Experiments were performed as in Fig. 5b.
e. DNA-bridging activity of FoxP3 orthologs. Experiments were performed as in Fig. 5c.
f. Sequence alignment of FoxP3 orthologs from different species, showing conservation of the key interface residues (yellow highlight, arrows on top with the residue identities in *M. musculus* FoxP3).
g. DNA-binding activity of FoxP3 paralogs with TnG repeats (n=1-4). Experiments were performed as in Fig. 5b.

**Extended data table 1**

ED table 1. Table for Cryo-EM data collection, refinement and validation statistics. The statistics of the 3D reconstruction and model refinement are summarized.

## Notes

### Competing Interest Statement

The authors have declared no competing interest.

### Summary of Updates

We have performed a series of experiments and analyses, and generated new data that resulted in 21 new figures and tables. These additional data further strengthen our manuscript.

## Main text references

1 Brunkow, M. E. et al. Disruption of a new forkhead/winged-helix protein, scurfin, results in the fatal lymphoproliferative disorder of the scurfy mouse. Nat Genet 27, 68–73, doi:10.1038/83784 (2001).

2 Bennett, C. L. et al. The immune dysregulation, polyendocrinopathy, enteropathy, X-linked syndrome (IPEX) is caused by mutations of FOXP3. Nat Genet 27, 20–21, doi:10.1038/83713 (2001).

3 Fontenot, J. D., Gavin, M. A. & Rudensky, A. Y. Foxp3 programs the development and function of CD4+CD25+ regulatory T cells. Nat Immunol 4, 330–336, doi:10.1038/ni904 (2003).

4 Hori, S., Nomura, T. & Sakaguchi, S. Control of regulatory T cell development by the transcription factor Foxp3. Science 299, 1057–1061, doi:10.1126/science.1079490 (2003).

5 Chatila, T. A. et al. JM2, encoding a fork head-related protein, is mutated in X-linked autoimmunity-allergic disregulation syndrome. J Clin Invest 106, R75–81, doi:10.1172/JCI11679 (2000).

6 Badis, G. et al. Diversity and complexity in DNA recognition by transcription factors. Science 324, 1720–1723, doi:10.1126/science.1162327 (2009).

7 Lambert, S. A. et al. The Human Transcription Factors. Cell 172, 650–665, doi:10.1016/j.cell.2018.01.029 (2018).

8 Inukai, S., Kock, K. H. & Bulyk, M. L. Transcription factor-DNA binding: beyond binding site motifs. Curr Opin Genet Dev 43, 110–119, doi:10.1016/j.gde.2017.02.007 (2017).

9 Kribelbauer, J. F., Rastogi, C., Bussemaker, H. J. & Mann, R. S. Low-Affinity Binding Sites and the Transcription Factor Specificity Paradox in Eukaryotes. Annu Rev Cell Dev Biol 35, 357–379, doi:10.1146/annurev-cellbio-100617-062719 (2019).

10 Reiter, F., Wienerroither, S. & Stark, A. Combinatorial function of transcription factors and cofactors. Curr Opin Genet Dev 43, 73–81, doi:10.1016/j.gde.2016.12.007 (2017).

11 Jolma, A. et al. DNA-dependent formation of transcription factor pairs alters their binding specificity. Nature 527, 384–388, doi:10.1038/nature15518 (2015).

12 Liu, Z., Lee, D. S., Liang, Y., Zheng, Y. & Dixon, J. Foxp3 Orchestrates Reorganization of Chromatin Architecture to Establish Regulatory T Cell Identity. bioRxiv, doi:10.1101/2023.02.22.529589 (2023).

13 Ramirez, R. N., Chowdhary, K., Leon, J., Mathis, D. & Benoist, C. FoxP3 associates with enhancer-promoter loops to regulate T(reg)-specific gene expression. Sci Immunol 7, eabj9836, doi:10.1126/sciimmunol.abj9836 (2022).

14 van der Veeken, J. et al. The Transcription Factor Foxp3 Shapes Regulatory T Cell Identity by Tuning the Activity of trans-Acting Intermediaries. Immunity 53, 971–984 e975, doi:10.1016/j.immuni.2020.10.010 (2020).

15 Zemmour, D. et al. Single-cell analysis of FOXP3 deficiencies in humans and mice unmasks intrinsic and extrinsic CD4(+) T cell perturbations. Nat Immunol 22, 607–619, doi:10.1038/s41590-021-00910-8 (2021).

16 Kwon, H.-K., Chen, H.-M., Mathis, D. & Benoist, C. Different molecular complexes that mediate transcriptional induction and repression by FoxP3. Nature Immunology 18, 1238–1248, doi:10.1038/ni.3835 (2017).

17 Hannenhalli, S. & Kaestner, K. H. The evolution of Fox genes and their role in development and disease. Nat Rev Genet 10, 233–240, doi:10.1038/nrg2523 (2009).

18 Benayoun, B. A., Caburet, S. & Veitia, R. A. Forkhead transcription factors: key players in health and disease. Trends Genet 27, 224–232, doi:10.1016/j.tig.2011.03.003 (2011).

19 Dai, S., Qu, L., Li, J. & Chen, Y. Toward a mechanistic understanding of DNA binding by forkhead transcription factors and its perturbation by pathogenic mutations. Nucleic Acids Res 49, 10235–10249, doi:10.1093/nar/gkab807 (2021).

20 Bandukwala, Hozefa S. et al. Structure of a Domain-Swapped FOXP3 Dimer on DNA and Its Function in Regulatory T Cells. Immunity 34, 479–491, doi:10.1016/j.immuni.2011.02.017 (2011).

21 Chen, Y. et al. DNA binding by FOXP3 domain-swapped dimer suggests mechanisms of long-range chromosomal interactions. Nucleic Acids Research 43, 1268–1282, doi:10.1093/nar/gku1373 (2015).

22 Leng, F. et al. The transcription factor FoxP3 can fold into two dimerization states with divergent implications for regulatory T cell function and immune homeostasis. Immunity 55, 1354–1369 e1358, doi:10.1016/j.immuni.2022.07.002 (2022).

23 Kitagawa, Y. et al. Guidance of regulatory T cell development by Satb1-dependent super-enhancer establishment. Nat Immunol 18, 173–183, doi:10.1038/ni.3646 (2017).

24 Samstein, R. M. et al. Foxp3 exploits a pre-existent enhancer landscape for regulatory T cell lineage specification. Cell 151, 153–166, doi:10.1016/j.cell.2012.06.053 (2012).

25 Jolma, A. et al. DNA-binding specificities of human transcription factors. Cell 152, 327–339, doi:10.1016/j.cell.2012.12.009 (2013).

26 Koh, K. P., Sundrud, M. S. & Rao, A. Domain requirements and sequence specificity of DNA binding for the forkhead transcription factor FOXP3. PLoS One 4, e8109, doi:10.1371/journal.pone.0008109 (2009).

27 Yoshida, H. et al. The cis-Regulatory Atlas of the Mouse Immune System. Cell 176, 897–912 e820, doi:10.1016/j.cell.2018.12.036 (2019).

28 Rubio-Cabezas, O. et al. Clinical heterogeneity in patients with FOXP3 mutations presenting with permanent neonatal diabetes. Diabetes Care 32, 111–116, doi:10.2337/dc08-1188 (2009).

29 Consonni, F., Ciullini Mannurita, S. & Gambineri, E. Atypical Presentations of IPEX: Expect the Unexpected. Front Pediatr 9, 643094, doi:10.3389/fped.2021.643094 (2021).

30 Ibrahim, A. et al. MeCP2 is a microsatellite binding protein that protects CA repeats from nucleosome invasion. Science 372, doi:10.1126/science.abd5581 (2021).

31 Contente, A., Dittmer, A., Koch, M. C., Roth, J. & Dobbelstein, M. A polymorphic microsatellite that mediates induction of PIG3 by p53. Nat Genet 30, 315–320, doi:10.1038/ng836 (2002).

32 Li, K., Luo, H., Huang, L., Luo, H. & Zhu, X. Microsatellite instability: a review of what the oncologist should know. Cancer Cell Int 20, 16, doi:10.1186/s12935-019-1091-8 (2020).

33 Pecina-Slaus, N., Kafka, A., Salamon, I. & Bukovac, A. Mismatch Repair Pathway, Genome Stability and Cancer. Front Mol Biosci 7, 122, doi:10.3389/fmolb.2020.00122 (2020).

34 Bonneville, R. et al. Landscape of Microsatellite Instability Across 39 Cancer Types. JCO Precis Oncol 2017, doi:10.1200/PO.17.00073 (2017).

35 Kloor, M. & von Knebel Doeberitz, M. The Immune Biology of Microsatellite-Unstable Cancer. Trends Cancer 2, 121–133, doi:10.1016/j.trecan.2016.02.004 (2016).

36 Bagshaw, A. T. M. Functional Mechanisms of Microsatellite DNA in Eukaryotic Genomes. Genome Biol Evol 9, 2428–2443, doi:10.1093/gbe/evx164 (2017).

37 Gharesouran, J., Hosseinzadeh, H., Ghafouri-Fard, S., Taheri, M. & Rezazadeh, M. STRs: Ancient Architectures of the Genome beyond the Sequence. J Mol Neurosci 71, 2441–2455, doi:10.1007/s12031-021-01850-6 (2021).

38 Braccioli, L. et al. FOXP1 Promotes Embryonic Neural Stem Cell Differentiation by Repressing Jagged1 Expression. Stem Cell Reports 9, 1530–1545, doi:10.1016/j.stemcr.2017.10.012 (2017).

39 Consortium, E. P. An integrated encyclopedia of DNA elements in the human genome. Nature 489, 57–74, doi:10.1038/nature11247 (2012).

## Additional references associated with Methods

40 Moparthi, L. & Koch, S. A uniform expression library for the exploration of FOX transcription factor biology. Differentiation 115, 30–36, doi:10.1016/j.diff.2020.08.002 (2020).

41 Dyer, P. N. et al. Reconstitution of nucleosome core particles from recombinant histones and DNA. Methods Enzymol 375, 23–44, doi:10.1016/s0076-6879(03)75002-2 (2004).

42 Punjani, A., Rubinstein, J. L., Fleet, D. J. & Brubaker, M. A. cryoSPARC: algorithms for rapid unsupervised cryo-EM structure determination. Nat Methods 14, 290–296, doi:10.1038/nmeth.4169 (2017).

43 Scheres, S. H. A Bayesian view on cryo-EM structure determination. J Mol Biol 415, 406–418, doi:10.1016/j.jmb.2011.11.010 (2012).

44 Kimanius, D., Dong, L., Sharov, G., Nakane, T. & Scheres, S. H. W. New tools for automated cryo-EM single-particle analysis in RELION-4.0. Biochem J 478, 4169–4185, doi:10.1042/BCJ20210708 (2021).

45 Zheng, S. Q. et al. MotionCor2: anisotropic correction of beam-induced motion for improved cryo-electron microscopy. Nat Methods 14, 331–332, doi:10.1038/nmeth.4193 (2017).

46 Rohou, A. & Grigorieff, N. CTFFIND4: Fast and accurate defocus estimation from electron micrographs. J Struct Biol 192, 216–221, doi:10.1016/j.jsb.2015.08.008 (2015).

47 Zivanov, J. et al. New tools for automated high-resolution cryo-EM structure determination in RELION-3. Elife 7, doi:10.7554/eLife.42166 (2018).

48 Pettersen, E. F. et al. UCSF Chimera--a visualization system for exploratory research and analysis. J Comput Chem 25, 1605–1612, doi:10.1002/jcc.20084 (2004).

49 Emsley, P., Lohkamp, B., Scott, W. G. & Cowtan, K. Features and development of Coot. Acta Crystallogr D Biol Crystallogr 66, 486–501, doi:10.1107/S0907444910007493 (2010).

50 Liebschner, D. et al. Macromolecular structure determination using X-rays, neutrons and electrons: recent developments in Phenix. Acta Crystallogr D Struct Biol 75, 861–877, doi:10.1107/S2059798319011471 (2019).

51 Williams, C. J. et al. MolProbity: More and better reference data for improved all-atom structure validation. Protein Sci 27, 293–315, doi:10.1002/pro.3330 (2018).

52 Morin, A. et al. Collaboration gets the most out of software. Elife 2, e01456, doi:10.7554/eLife.01456 (2013).

53 Ohi, M., Li, Y., Cheng, Y. & Walz, T. Negative Staining and Image Classification -Powerful Tools in Modern Electron Microscopy. Biol Proced Online 6, 23–34, doi:10.1251/bpo70 (2004).

54 Langmead, B. & Salzberg, S. L. Fast gapped-read alignment with Bowtie 2. Nat Methods 9, 357–359, doi:10.1038/nmeth.1923 (2012).

55 Danecek, P. et al. Twelve years of SAMtools and BCFtools. Gigascience 10, doi:10.1093/gigascience/giab008 (2021).

56 Zhang, Y. et al. Model-based analysis of ChIP-Seq (MACS). Genome Biol 9, R137, doi:10.1186/gb-2008-9-9-r137 (2008).

57 Bailey, T. L., Johnson, J., Grant, C. E. & Noble, W. S. The MEME Suite. Nucleic Acids Res 43, W39–49, doi:10.1093/nar/gkv416 (2015).

58 Bailey, T. L. STREME: accurate and versatile sequence motif discovery. Bioinformatics 37, 2834–2840, doi:10.1093/bioinformatics/btab203 (2021).

59 Quinlan, A. R. & Hall, I. M. BEDTools: a flexible suite of utilities for comparing genomic features. Bioinformatics 26, 841–842, doi:10.1093/bioinformatics/btq033 (2010).

60 Grant, C. E., Bailey, T. L. & Noble, W. S. FIMO: scanning for occurrences of a given motif. Bioinformatics 27, 1017–1018, doi:10.1093/bioinformatics/btr064 (2011).

61 Yu, G., Wang, L. G. & He, Q. Y. ChIPseeker: an R/Bioconductor package for ChIP peak annotation, comparison and visualization. Bioinformatics 31, 2382–2383, doi:10.1093/bioinformatics/btv145 (2015).

62 Ramirez, F. et al. deepTools2: a next generation web server for deep-sequencing data analysis. Nucleic Acids Res 44, W160–165, doi:10.1093/nar/gkw257 (2016).

63 Kwon, H. K., Chen, H. M., Mathis, D. & Benoist, C. Different molecular complexes that mediate transcriptional induction and repression by FoxP3. Nat Immunol 18, 1238–1248, doi:10.1038/ni.3835 (2017).

