## Supplementary figures and images for "FoxP3 recognizes microsatellites and bridges DNA through multimerization"

### Extended Data Figure 1

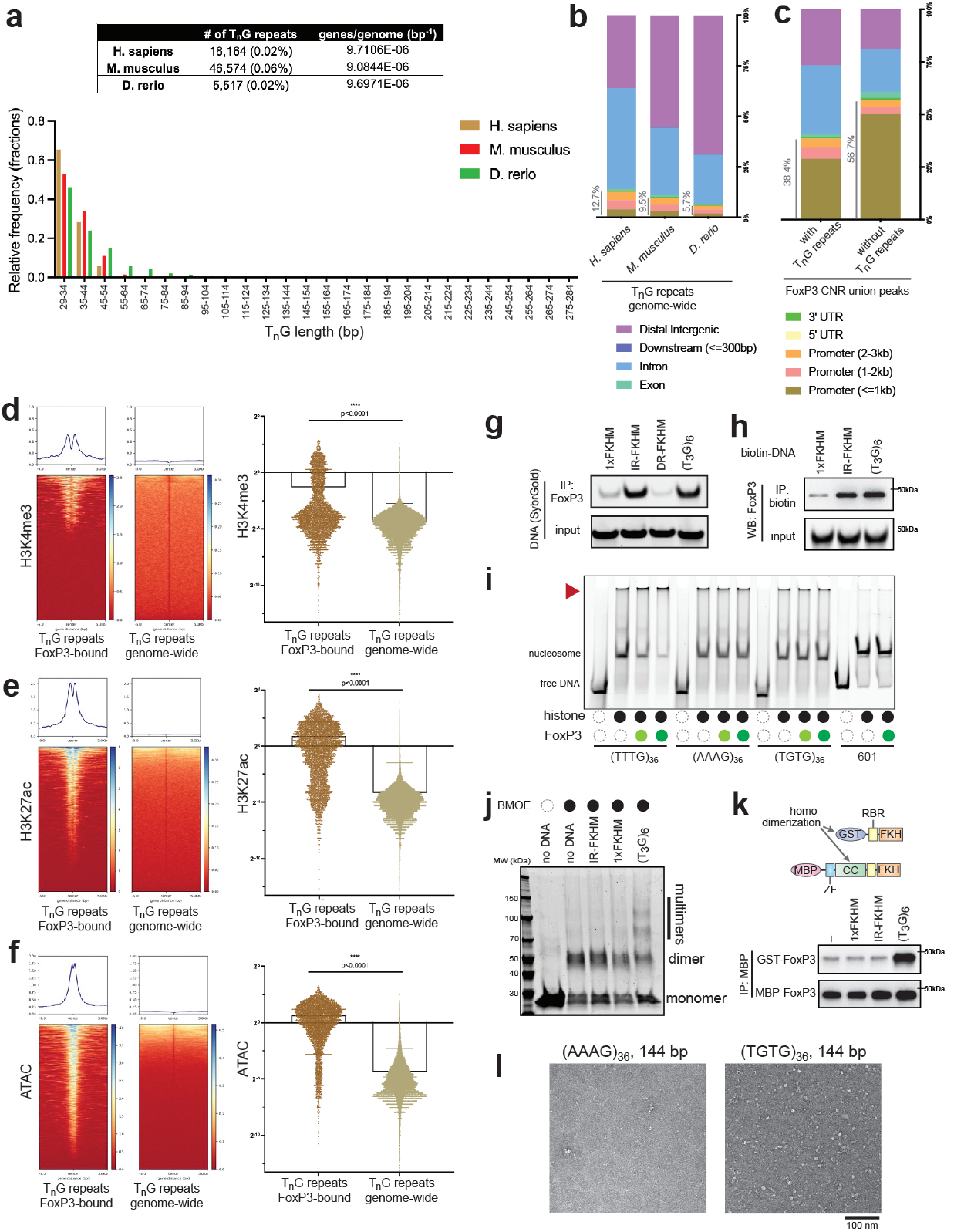

### Extended Data Figure 2

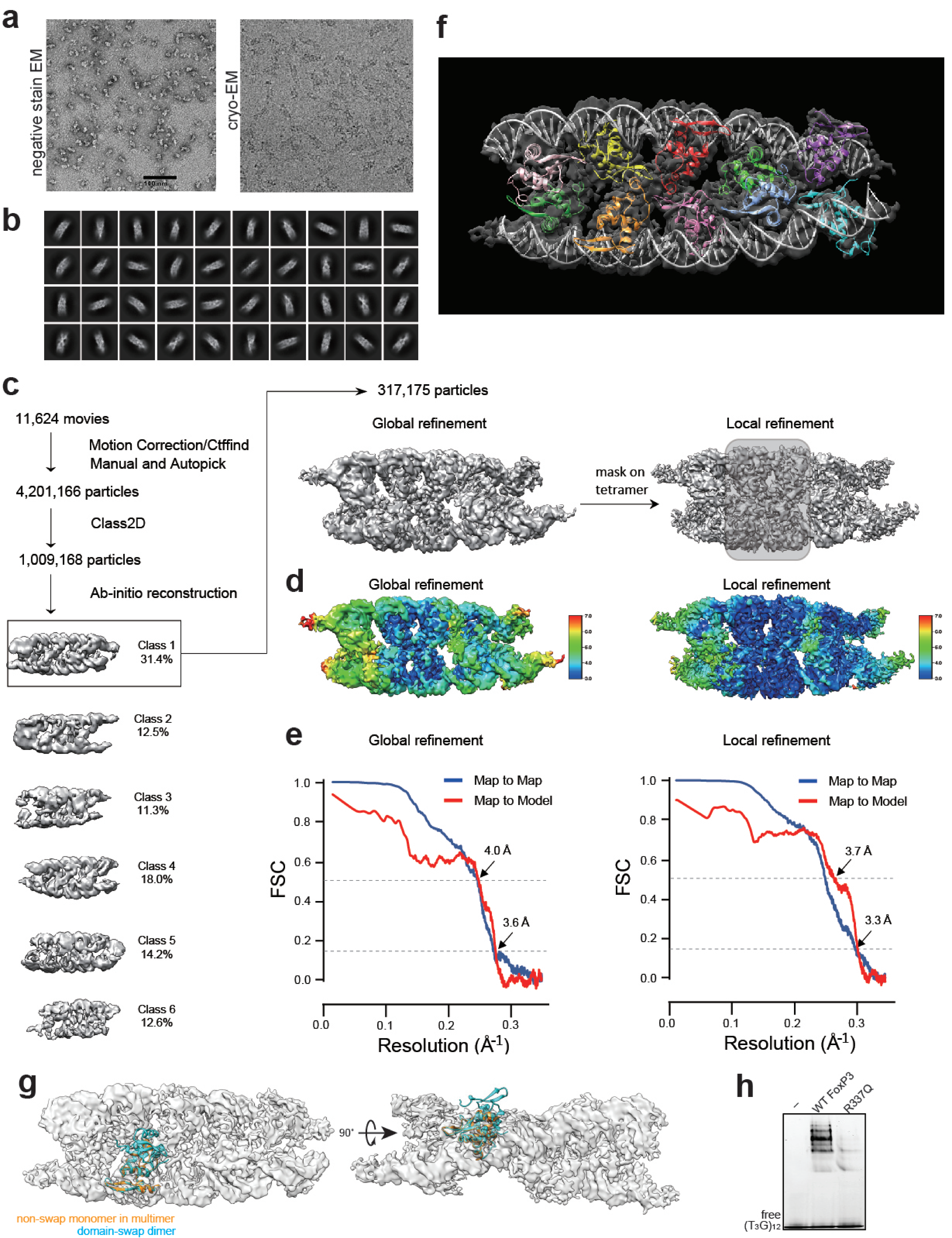

### Extended Data Figure 3

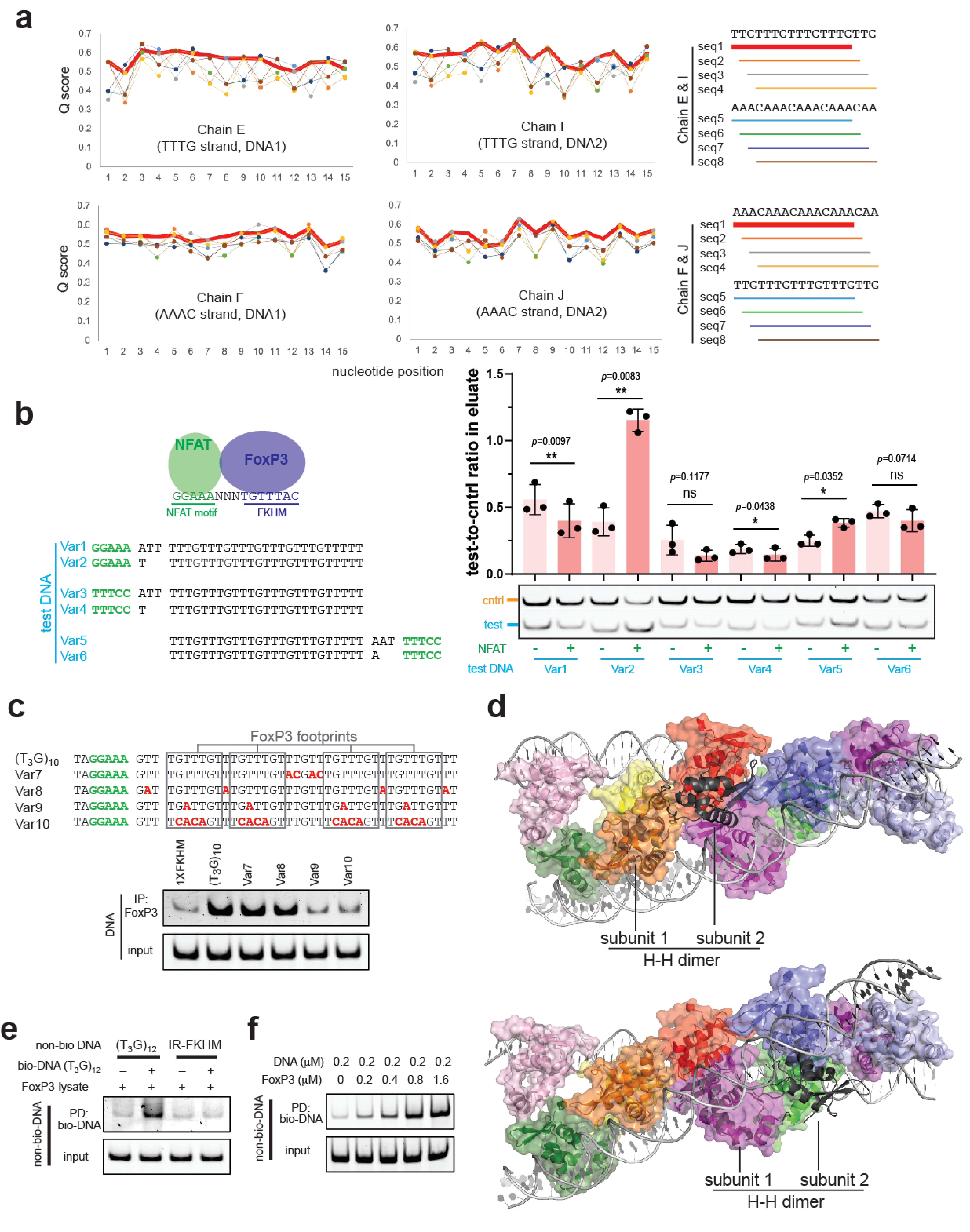

### Extended Data Figure 4

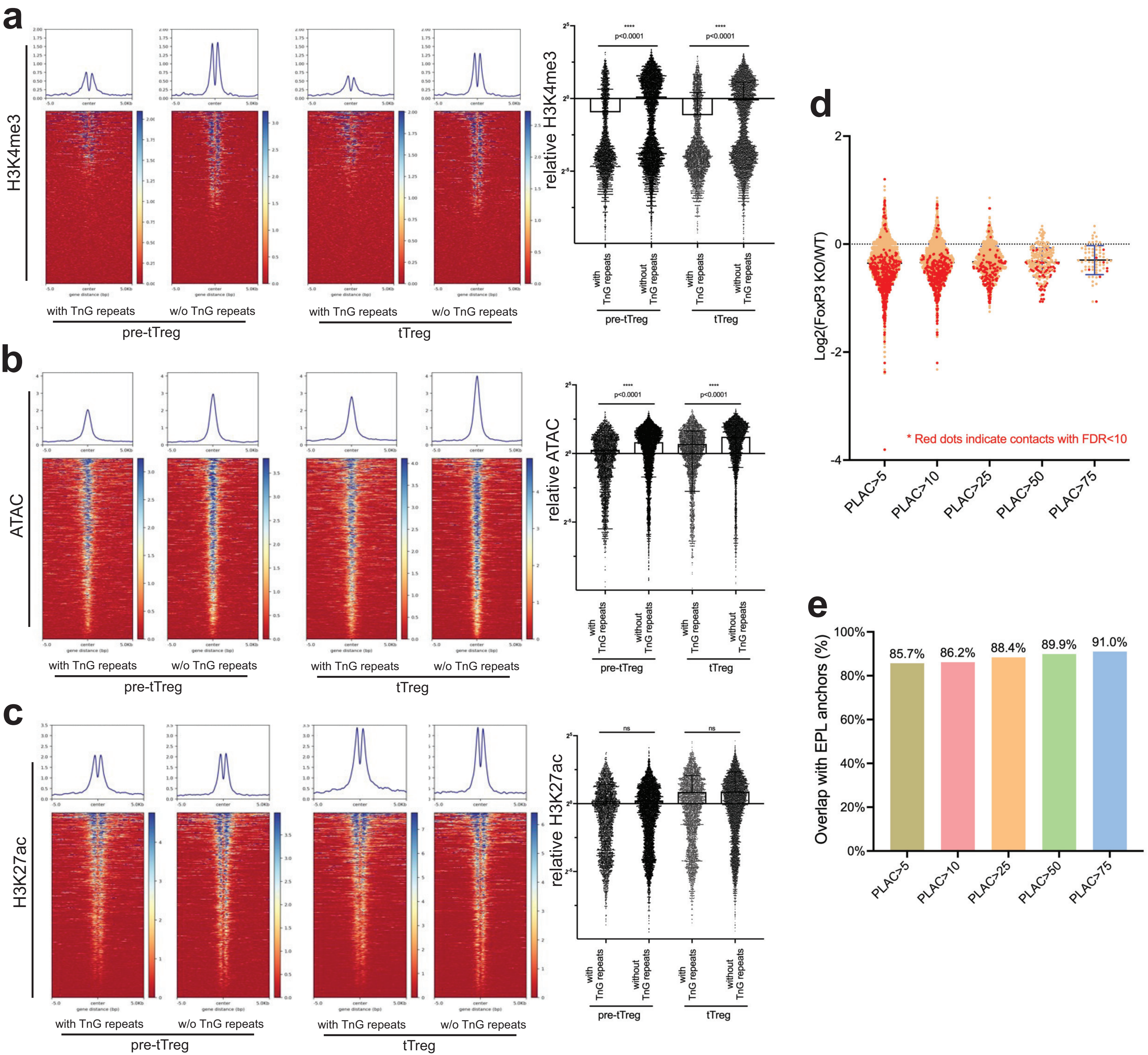

### Extended Data Figure 5

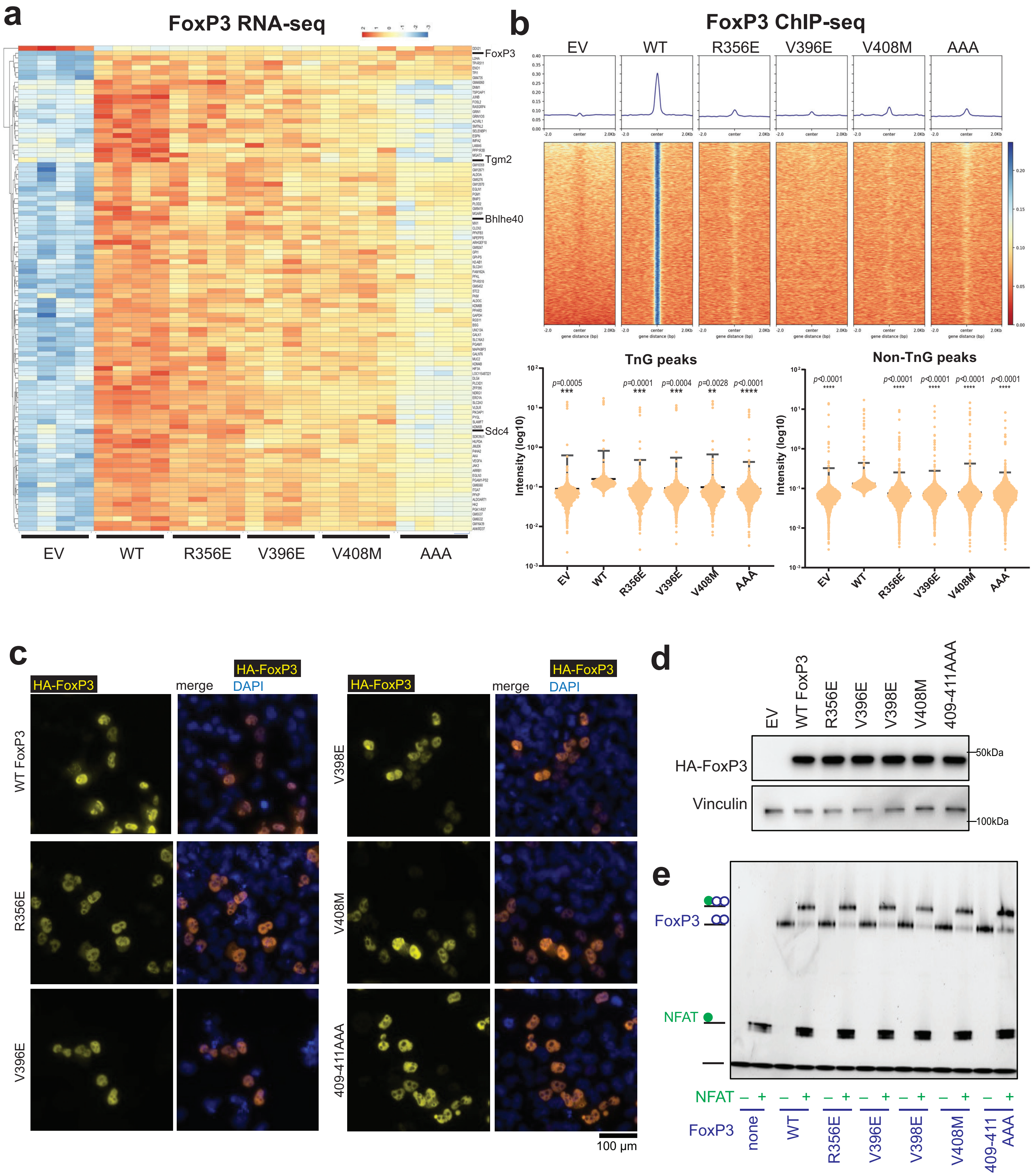

### Extended Data Figure 6

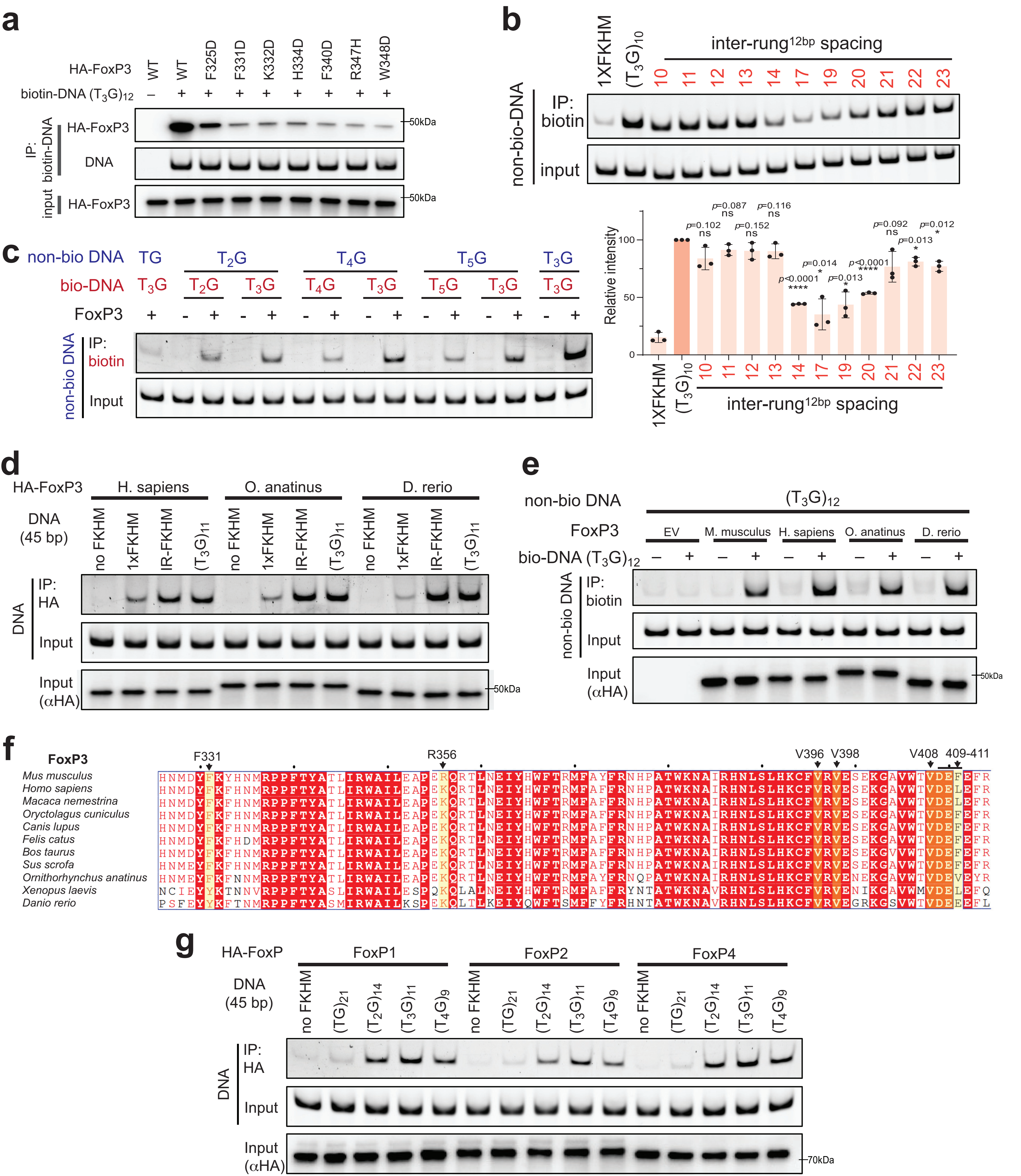
