## Extended Data Table 1 for "FoxP3 recognizes microsatellites and bridges DNA through multimerization"

**Cryo-EM data collection, refinement and validation statistics**

|  | #1 Decamer of FOXP3^ΔN^-TTTG18  (EMDB-40737)  (PDB 8SRP) | #2 Tetramer of FOXP3^ΔN^-TTTG18  (EMDB-40736)  (PDB 8SRO) |
| --- | --- | --- |
| **Data collection and processing** |  |  |
| Magnification | 81000 | 81000 |
| Voltage (kV) | 300 | 300 |
| Electron exposure (e–/Å^2^) | 60 | 60 |
| Defocus range (μm) | -0.7 to -2.1 | -0.7 to -2.1 |
| Pixel size (Å) | 0.844 | 0.844 |
| Symmetry imposed | C1 | C1 |
| Initial particle images (no.) | 4201166 | 4201166 |
| Final particle images (no.) | 317175 | 317175 |
| Map resolution (Å)  FSC threshold | 3.6  0.143 | 3.3  0.143 |
| **Refinement** |  |  |
| Initial model used (PDB code) | 7TDX | 7TDX |
| Model resolution (Å)  FSC threshold | 4.0  0.5 | 3.7  0.5 |
| Map sharpening *B* factor (Å^2^) | -168.5 | -161 |
| Model composition  Non-hydrogen atoms  Protein residues  Ligands | 9516  849  214 | 4132  345  72 |
| R.m.s. deviations  Bond lengths (Å)  Bond angles (°) | 0.004  0.658 | 0.003  0.559 |
| Validation  Clashscore  Poor rotamers (%) | 9.34  0.47 | 8.41  0.90 |
| Ramachandran plot  Favored (%)  Allowed (%)  Disallowed (%) | 93.49  6.51  0 | 94.66  5.34  0 |
